# The direct and indirect effects of environmental toxicants on the health of bumble bees and their microbiomes

**DOI:** 10.1101/2020.04.24.060228

**Authors:** Jason A. Rothman, Kaleigh A. Russell, Laura Leger, Quinn S. McFrederick, Peter Graystock

## Abstract

Bumble bees (*Bombus* spp.) are important and widespread insect pollinators, but the act of foraging on flowers can expose them to harmful pesticides and environmental chemicals such as oxidizers and heavy metals. How these compounds directly influence bee survival and indirectly affect bee health via the gut microbiome is largely unknown. As the toxins and toxicants in floral nectar and pollen take many forms, we explored the genomes of core bumblebee microbes (Using RAST) for their potential to detoxify cadmium, copper, selenium, the neonicotinoid pesticide imidacloprid, and hydrogen peroxide - which have all been identified in floral nectar and pollen. We then exposed *Bombus impatiens* workers to varying concentrations of these chemicals spiked into their diet and identified the direct effects on bee survival. Using field realistic doses, we further explored indirect effects on bee microbiomes. We found multiple genes in core gut microbes that have the potential to aid in detoxifying harmful chemicals. We also found that while the chemicals are largely toxic at levels within and above field-realistic concentrations, the field-realistic concentrations - except for imidacloprid - altered the composition of the bee microbiome, potentially causing gut dysbiosis. Overall, our study shows that environmental chemicals found in floral nectar and pollen can directly cause bee mortality, and at field-realistic levels, likely have indirect, deleterious effects on bee health via their influence on the bee microbiome.

## Introduction

Wild and managed bumble bees (*Bombus* spp.) are globally valuable pollinators for both crops and wild flowering plants (1,2). It is therefore worrisome that some populations of these important pollinators are declining in both Europe (3) and North America (4), with the main contributors of these declines thought to be habitat destruction, disease, and chemical exposure (5,6). While pesticides are considered to be the primary class of xenobiotics responsible for bee decline, foraging bees are exposed to additional environmental toxicants in floral nectar and pollen such as heavy metals (7), and oxidizers (8). The direct risks these chemicals pose to bees is not fully appreciated, furthermore, the indirect influences most chemicals (including pesticides) have on bee health is an understudied area (9,10).

Chemical exposure occurs primarily to bees while foraging on flowers. Flowers provide essential nutrition; however, floral nectar and pollen can also harbor chemicals toxic to bees. Research on the chemical toxicology of plants has focused heavily on the presence of pesticides such as neonicotinoids in floral nectar and pollen (11). In addition to pesticides, floral pollen and nectar may contain naturally produced oxidizers such as hydrogen peroxide, plus heavy metals sequestered from contaminated soils (12–15). Flowers produce oxidizers (16) and sequester heavy metals (17) to protect themselves from parasites and herbivores respectively. The hive products of honey bees foraging on flowers that are growing in soils contaminated with selenium ions, copper, and/or cadmium have been shown to accumulate these pollutants at concentrations greater than in the flowers (7). So frequent is this exposure and accumulation of contaminants that honey bees have been proposed as bioindicators of soil contamination whereby testing the hives’ honey for metals will indicate the presence of local environmental toxins (18).

The exposure of bees to xenobiotics identified in floral products may contribute to the declines in solitary and social bees alike. Honey bees, bumble bees (*Bombus* spp.) and mason bees (*Osmia* spp.), all rapidly die after exposure to high concentrations of heavy metals and pesticides in lab experiments (19–21). Field-realistic doses of pesticides in flowers can also lead to reduced bee survival and reductions in hive fitness (5). It is unknown what effects field-realistic doses of heavy metals or oxidizers such as hydrogen peroxide may have on foraging bees, particularly if consumed over a period of days as would naturally occur. In addition to this direct lethality, increasing evidence suggests that exposure to low doses can cause sublethal damage to bee health (22). Sublethal effects on bee health are known to often have cascading and negative impacts on the overall vigor of bee colonies and populations, and consequently are of significant importance.

The gut microbiome is considered to be a crucial factor in bee health (23). The distinct gut microbiomes present in social bees are largely consistent worldwide (24–28) and positively affect bee health via toxicant metabolism (29), immune system stimulation (30) and protection against pathogens (25,31,32). While the bacterial members of the microbiome remain generally consistent, stressors have been found to alter the microbial community, as found during times of inconsistent forage availability (33,34), antibiotic exposure (35), infection (36), and pesticide exposure (37). During times of host stress, the symbiotic community of bacteria within the gut can collapse, removing its positive effect on bee health and effectively causing dysbiosis of the gut microbiome. Such dysbiosis may further reduce bee health, making the bees vulnerable to nutritional stress and disease (38–40).

The effects of xenobiotic chemicals on the microbiomes of animals is emerging as an integral part of the ecotoxicology of harmful compounds (41). Cadmium (42,43), selenium (44), copper (45), imidacloprid (46), and hydrogen peroxide (as measured as ROS (47)) have been shown to alter host microbiomes in non-bee study systems. Outside of our work, little is known about how these toxicants affect bee microbiota, with field-realistic selenium and cadmium exposure altering proportions of microbes in honey bees (22), and imidacloprid not affecting the microbiome of honey bees (48). While the effects of toxicants on host-associated microbes have been studied in many systems, a new paradigm is emerging in which the microbiome potentially protects its host from metal(loid) toxicity. For example, host-associated bacteria have been shown to detoxify chromium and lead (49), copper (50), arsenic (51), and selenite (52), mainly through accumulation or respiration. Specifically in bees, the microbiome has been shown to reduce selenate-induced mortality (53), and bee-associated microbes removed cadmium from their environment (22), although the mechanisms for microbe-mediated protection against toxicants in bees remains largely unknown. As bacterial genes often encode metal(loid) transporters and detoxification pathways, we may be able to understand toxicant-protection mechanisms by annotating the pathways in symbiont genomes.

Given the importance of environmental toxicants and the microbiome in host health, we investigated the interactions between multiple chemical poisons, the bumble bee *B. impatiens*, and its associated microbes. First, we searched the genomes of bee associated microbes for evidence they could play a role in the metabolism/detoxification of common chemicals (selenate, copper, cadmium, imidacloprid and hydrogen peroxide). Second: we tested the direct lethality of these toxicants to bumble bees. Third: we determined if exposure to natural concentrations of selenate, copper, cadmium, imidacloprid or hydrogen peroxide altered the healthy microbiome of the bee.

## Materials and Methods

### Identifying evidence of microbial transport or detoxification via genome annotations

To identify the genomic basis for toxicant tolerance/transport we annotated publicly available genomes of bee symbionts with the RAST Server (Rapid Annotations using Subsystems Technology) (54) using whole genome sequence data obtained from the National Center for Biotechnology Information (NCBI). Based on known “core” microbes and opportunistic microbes commonly found within the microbiomes of bumble bees (27), we annotated genomes from strains of the following species: *Bifidobacterium bombi*, *Bifidobacterium commune*, *Bombella intestini*, *Bombiscardovia coagulans*, Candidatus *Schmidhempelia bombi*, *Commensalibacter intestini*, *Gilliamella apicola*, *Lactobacillus bombicola*, *Serratia marcescens*, and *Snodgrassella alvi* (see Supplemental file SF1 for strain IDs and genome accession numbers). We used the following RAST subsystems to narrow our searches: “Cobalt-zinc-cadmium resistance”, “copper homeostasis”, “copper homeostasis copper tolerance”, “copper transport system”, “oxidative stress tolerance”, “selenate/selenite uptake”, and “selenocysteine metabolism.”

### Bumble bee rearing and toxicity tests for each compound

We purchased 10 commercial bumble bee (*Bombus impatiens*) colonies from Koppert Biological Systems, Inc. (Howell, MI) that contained a mated queen, ~200 workers, pollen, and proprietary sugar solution. We immediately replaced the proprietary sugar solution with 60% sucrose and provided the colonies with pollen patties ad libitum. To allow the colonies to develop, we kept them under constant darkness at 29°C in temperature-controlled rooms at the University of California, Riverside for two weeks before starting the experiment. We collected 60 adult worker bees from each of three colonies (N = 180 bees for each of the five compounds) and sorted them by colony into cohorts of five bees in 475mL polypropylene containers (WebstaurantStore, Lancaster, PA). Based on published ranges (Supplemental table ST1), we exposed the bees to the following treatments: 10 mg/L, 1.0 mg/L, 0.1 mg/L, 0.01 mg/L, 0.001 mg/L, and 0 mg/L spiked into 60% sucrose for sodium selenate, cadmium chloride, and imidacloprid, 100 mg/L, 10 mg/L, 1.0 mg/L, 0.1 mg/L, 0.01 mg/L and 0 mg/L copper chloride spiked into 60% sucrose, and 1.0 M, 0.1 M, 0.01 M, 0.001 M, 0.0001 M, and 0 M hydrogen peroxide spiked into 60% sucrose. We allowed the bees to feed *ad libitum* for 14 days while we recorded mortality daily. To analyze survivorship, we used the R packages “drc,” (55) to calculate three-parameter log-logistic functions for model picking, and “survival” (56) to calculate statistical significance, hazard models, and test our data to ensure that it met proportional hazards assumptions, and “survminer” to visualize the survival curves (57).

### Indirect effects of toxicity as observed on the microbiome

We purchased three additional bumble bee colonies from Koppert Biological Systems, Inc. and reared the bees in the same manner as above. To expose bees to toxicants, we isolated 60 mature workers from each colony (N = 180) in 60 mL polypropylene containers (WebstaurantStore, Lancaster, PA). We exposed bees to the chemical treatments by chronically feeding 30 bees 60% sucrose spiked with either 0.25 mg/L cadmium chloride (Sigma Aldrich, St. Louis, MO), 0.5 mg/L sodium selenate (Alfa Aesar, Ward Hill, MA), 25 mg/L copper chloride (Sigma Aldrich, St. Louis, MO), 0.001 mg/L imidacloprid (Sigma Aldrich, St. Louis, MO), 0.025 mol/L hydrogen peroxide (Fisher Scientific, Waltham, MA), or 60% sucrose as a control (N = 30 per treatment), based on concentrations within the dose response assay and published ranges (Supplemental table ST1). We allowed the bees to feed on either toxicant-spiked or control sucrose ad libitum for four days and then flash froze the bees and stored them at −80°C.

### DNA extractions and 16S rRNA gene sequencing library preparation

We used a DNA extraction protocol based on Engel *et al* 2013 (58), Pennington *et al* 2017 (59), and Rothman *et al* 2019 (60). We first surface sterilized individual bees using a 0.1% sodium hypochlorite wash followed by three rinses with ultrapure water. We then sterilely dissected the whole gut out of each bee and transferred the gut into DNeasy Blood and Tissue Kit lysis plates (Qiagen, Valencia, CA) containing approximately 100 μL of 0.1mm glass beads, one 3.4mm steel-chrome bead (Biospec, Bartlesville, OK) and 180 μL of buffer ATL, followed by homogenization with a Qiagen Tissuelyser at 30 Hz for 6 minutes. We followed the remainder of the Qiagen DNeasy Blood and Tissue Kit protocol after homogenization. We also included four blanks to control for reagent contamination, which we prepared and sequenced in the same way as samples.

We prepared 16S rRNA gene libraries for paired-end Illumina MiSeq sequencing for each individual bee (N=134) using the protocol from McFrederick and Rehan 2016 (61), Pennington et al. 2018 (62) and Rothman et al. 2018 (33). We incorporated the 16S rRNA gene primer sequence, unique barcode sequence, and Illumina adapter sequence as in (63). We used the primers 799F-mod3 (64) and 1115R (63) for PCR with the following reaction conditions : 4 μL of template DNA, 0.5 μL of 10μM 799F-mod3, 0.5 μL of 10μM 1115R, 10 μL PCR grade water and 10 μL Pfusion DNA polymerase (New England Biolabs, Ipswich, MA), at an annealing temperature of 52°C for 30 cycles in a C1000 Touch thermal cycler (BioRad, Hercules, CA). We then removed excess primers and dNTPs with a PureLink Pro 96 PCR Purification Kit (Invitrogen, Carlsbad, CA) and used the cleaned PCR amplicons as a template for another PCR with the primers PCR2F and PCR2R to complete the Illumina adapter sequence (63). We performed PCR with the following reaction conditions: 0.5 μL of 10μM forward primer, 0.5 μL of 10μM reverse primer, 1 μL of template, 13 μL of ultrapure water and 10 μL of Pfusion DNA polymerase at an annealing temperature of 58°C for 15 cycles. We normalized the resulting libraries with a SequalPrep Normalization kit by following the supplied protocol (ThermoFisher Scientific, Waltham, MA). We pooled 5 μL of each normalized library and performed a final clean up with a single column PureLink PCR Purification Kit, then sequenced the multiplexed libraries using a V3 Reagent Kit at 2 × 300 cycles on an Illumina MiSeq Sequencer in the UC Riverside Genomics Core Facility. Raw sequencing data are available on the NCBI Sequence Read Archive (SRA) under accession numbers SRR6788889 - SRR6789022, and microbiome data of selenate versus control treatments were previously published in Rothman et al. 2019 (53).

### Microbiome bioinformatics and statistics

We used QIME2-2018.6 (65) to process the 16S rRNA gene sequence libraries. First, we trimmed the low-quality ends off the reads with QIIME2 and then used DADA2 (66) to bin our sequences into exact sequence variants (ESVs; 16S rRNA gene sequences that are 100% matches), remove chimeras, and remove reads with more than two expected errors. We assigned ESV taxonomy using the q2-feature-classifer (67) trained to the 799-1115 region of the 16S rRNA gene with the SILVA database (68). We also conducted BLASTn searches against the NCBI 16S microbial database (July 2018). We filtered out ESVs from the resulting table that corresponded to reagent contaminants as identified in our blanks or were assigned as chloroplast or mitochondria. We used the MAFFT aligner (69) and FastTree v2.1.3 to generate a phylogenetic tree of our sequences (70). We then used this tree and ESV table (Supplemental File SF2) to analyze alpha diversity and to tabulate UniFrac distance matrices. We visualized the UniFrac distances through two-dimensional Principal Coordinates Analysis (PCoA) with the R package “ggplot2” (71). We analyzed the alpha diversity of our samples through the Shannon Diversity Index and the Kruskal-Wallis test in QIIME2. Lastly, we tested our beta diversity data for statistical significance in R v3.5.1 (72) with the packages “vegan” (73) and “DESeq2” (74). Data and representative code can be found on Data Dryad (doi.org/10.7280/D14T2K) upon publication.

## Results

### Genomic bases of chemical resistance

Through our RAST annotations of core microbial genomes, we identified several genes that suggest that microbial genera commonly associated with bumble bees could reduce the negative effects of these toxic compounds to bee health. Several bee symbionts and other bacteria identified by our next-generation sequencing study had some or all of the following genes annotated in their genomes (Fig. S1). For selenium ion resistance, we found genes corresponding to the selenium ion transporters DedA (75), TsgA (76), and putative selenium ion and sulfate importer CysA (77). For cadmium ion resistance, we found the genes CzcABC, which encode the components of a cation transporter (78), its response regulator CzcD (79), and a cadmium-responsive transcriptional regulator, CadR (80). We identified the following genes involved in copper resistance: A copper-translocating ATPase (81), two copper-binding multicopper oxidases (82,83) (SufI and CueO, respectively), genes encoding the likely copper-binding proteins ScsABCD and CutEF (84), components of a copper-sequestering protein complex CopCD (85), and a copper-responsive transcriptional regulator, CueR (86). Lastly, we searched for genes involved in oxidative stress response and found genes encoding paraquat-inducible superoxide dismutase (SOD) PqiAB, Mn- and Fe-SODs (87), the SOD response regulon SoxS (88), a LysR-family peroxide-inducible transcriptional regulator (89), ferroxidase, a ferric uptake regulation protein (FUR) (90), the zinc/copper uptake regulation protein Zur, which may protect against oxidative stress (91), the antioxidant gene NnrS (92), an Fnr-like transcriptional regulator (93), catalase/peroxidase (94), and alkyl hyperoxide reductase C (AhpC) (95).

As *S. alvi* and *G. apicola* genomes are known to vary between strains (96,97) and there are several genomes for each taxon publicly available, we compared the above-mentioned detoxification/tolerance genes across strains within these species (53 strains of *S. alvi* and 67 strains of *G. apicola*). We found that *G. apicola* had notable variation across genes involved in responding to oxidative stress (specifically NnrS, SoxS, Fnr, and catalase), copper tolerance (the copper-translocating ATPase and SufI), cadmium tolerance (CadR), and overall selenate tolerance. There was less overall variation in detoxification/tolerance genes across *S. alvi* strains: we found strain variation in copper (CueR, CueO, and the copper-translocating ATPase) and cadmium tolerance (CzcA and CadR), while there was no genetic variation in oxidative stress response or selenate tolerance (Fig. 1).

**Figure 1.**
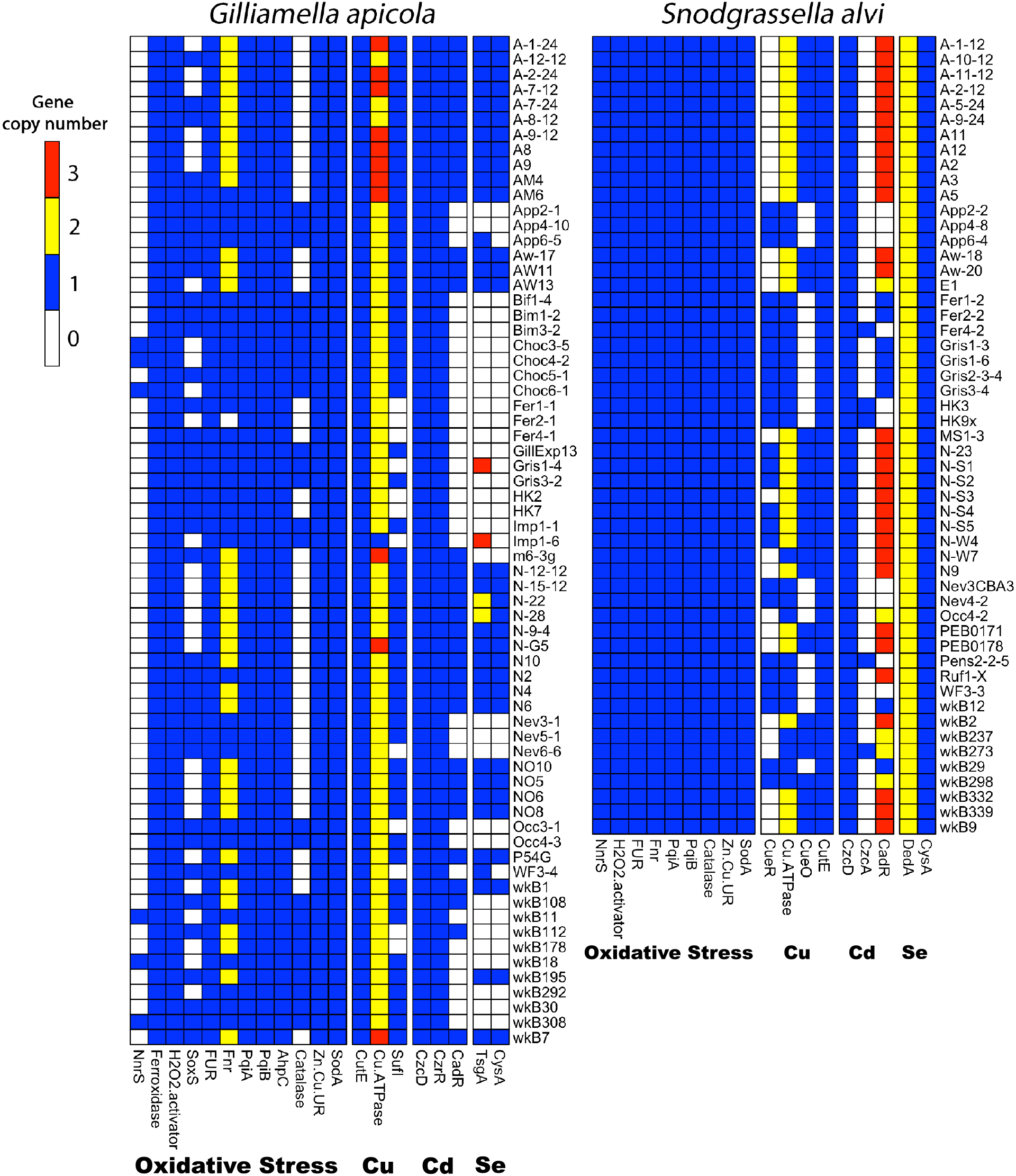
Illustration of the toxicant-tolerance genes found in strains of *Snodgrassella alvi* and *Gilliamella apicola* as annotated by RAST. Colored cells represent the copy number of each gene, row names indicate the bacterial strain, and column names denote the gene abbreviation or name and genes are grouped by type of stress. Nonstandard gene abbreviations are as follows: “H_2_O_2_.activator” is a peroxide-inducible genes activator, “FUR” is a ferric uptake regulation protein, “Fnr” is a fumarate and nitrate reduction regulatory protein, “Zn.Cu.UR” is a zinc/copper uptake regulation protein, and “Cu.ATPase” is a copper-translocating ATPase. natural range, all other chemicals were within natural ranges.

### Direct toxicity of each compound on bumble bee survival

We found that the concentration ranges of cadmium, copper, selenate, imidacloprid, and hydrogen peroxide went from no deaths (zero indirect effects on survival) to complete mortality. Over seven days of continuous exposure, survival in the various concentrations differed significantly (cox proportional hazard test log rank P < 0.001 for each compound, Fig. 2 and Fig. S2). We note that the lowest concentrations did not affect survival, and that concentrations above mg/L cadmium, 100 mg/L copper, 1 mg/L selenate, 0.1 mg/L imidacloprid, and 1.0 M hydrogen peroxide significantly reduced bee survival, which indicated a dose-dependent response (Supplemental Table ST2). We also calculated the LC_50_ after seven days continuous exposure for each toxicant: cadmium: 0.83 mg/L, copper: 66.55 mg/L, imidacloprid: 0.22 mg/L, selenate: 0.75 mg/L, and hydrogen peroxide: 0.39 mol/L (Fig. S3). We note that while we exposed bees to treatments for 14 days, the data from copper and cadmium did not fit the proportional hazards assumptions due to high mortality in controls after 11 and 9 days respectively. Survival results of these compounds up to seven days fit the assumptions, so all five treatments were tested over a total of seven days. Additionally, we tested selenate, imidacloprid, and hydrogen peroxide for 14 days and report the 14-day LC_50_ as follows: selenate: 0.09 mg/L, imidacloprid: 0.11 mg/L, and hydrogen peroxide: 0.27 mol/L. (Fig. S3 & S4).

**Figure 2.**
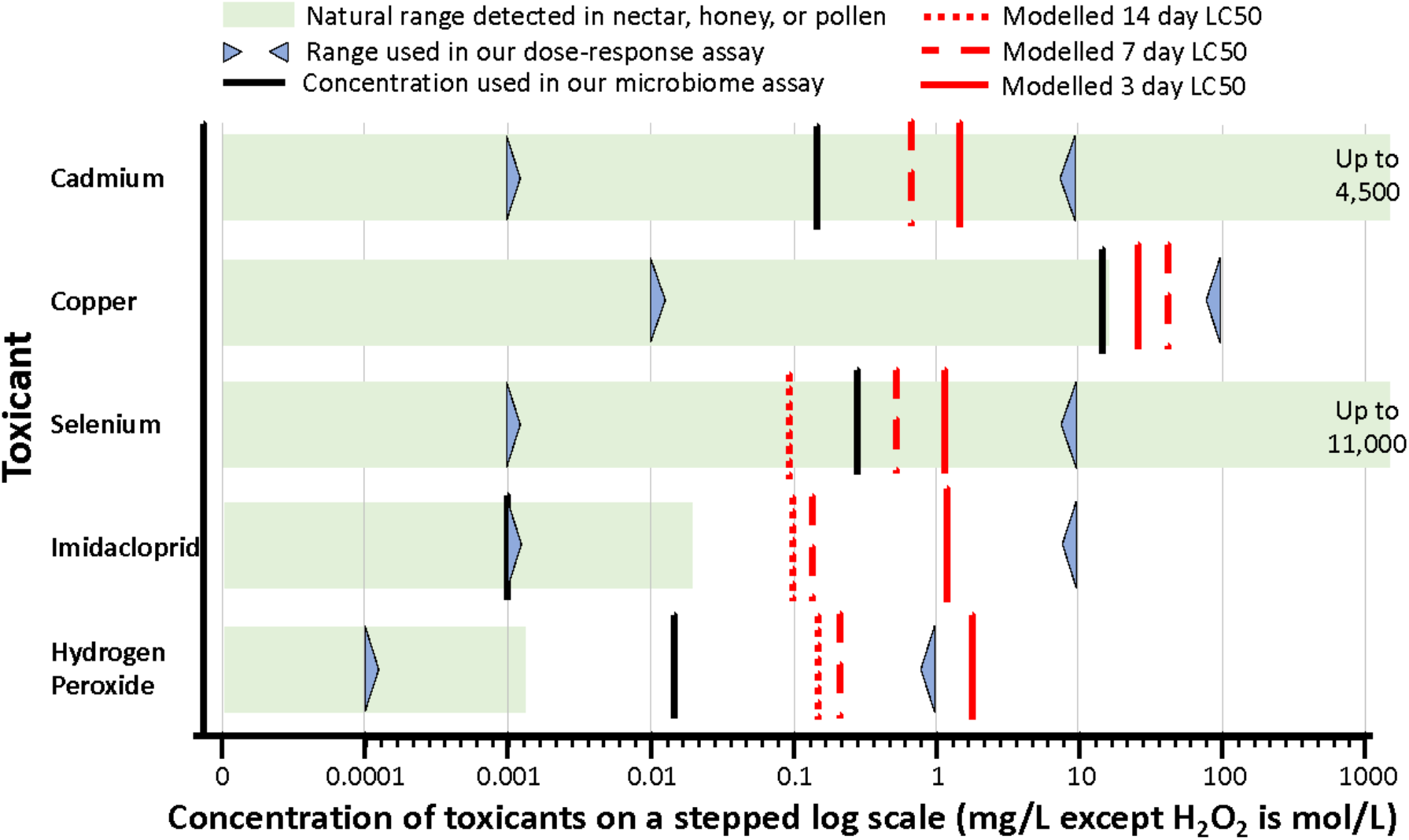
Natural ranges reported in nectar, honey, or pollen for each cadmium, copper, imidacloprid, hydrogen peroxide, and selenate along with the concentration ranges used in our dose-response and microbiome assays. Also annotated are the cox proportional hazard modelled LC50 values for each toxicant after three days, seven days, and ten days of continuous exposure. Note in the microbiome assay, the concentration of H_2_O_2_ used was ten times greater than the natural range.

### Amplicon sequencing alpha diversity and library statistics

We obtained 743,529 quality-filtered 16S rRNA gene sequences with a mean of 5,467 reads per sample (N = 136) that clustered into 113 Exact Sequence Variants (ESVs; sequences that are 100% identical). We determined that our samples had a representative coverage of bacterial diversity at a sequencing depth of 2,182 reads through rarefaction analysis (Fig. S5), as the curves saturated at approximately 1,110 reads. Overall, alpha diversity was significantly different due to treatment (Shannon’s H = 24.21, P < 0.001), although pairwise Kruskal-Wallis tests indicated that only selenate treatments had higher diversity as compared to controls (Benjamini-Hochberg corrected P_adj_ < 0.05, Supplemental Table ST3).

### Beta diversity and differential abundance of bacterial taxa between treatments

Regardless of treatment, we found that the gut communities of our samples were composed of bacteria of the genera *Gilliamella, Snodgrassella, Lactobacillus, Bifidobacterium, Bombiscardovia, Commensalibacter,* and *Serratia,* while other bacteria accounted for less than 1% of the relative abundance. To visualize the bumble bee gut microbiota, we generated a stacked bar plot representing bacteria present in greater than 1% proportional relative abundance in each sample (Fig. 3 and Fig. S6) and beta-diversity through Principal Coordinates Analysis (PCoA, Fig. 4), with only copper clearly clustering separately from control. We analyzed the Generalized UniFrac distance matrix of our samples with Adonis PERMANOVA (999 permutations) using colony and treatment as covariates and found that overall, there was a significant effect of treatment (F = 4.57, R^2^ = 0.14, P < 0.001), colony (F = 6.71, R^2^ = 0.08, P < 0.001) and interaction of these factors (treatment X colony, F = 1.63, R^2^ = 0.10, P < 0.001). As we had multiple separate treatments, we analyzed the pairwise interactions between each treatment versus control and found that each treatment except imidacloprid caused a significant change to the beta diversity of the bees’ microbiomes (Benjamini-Hochberg corrected for each treatment P_adj_ < 0.02; imidacloprid: P_adj_ = 0.96, Table ST3).

**Figure 3.**
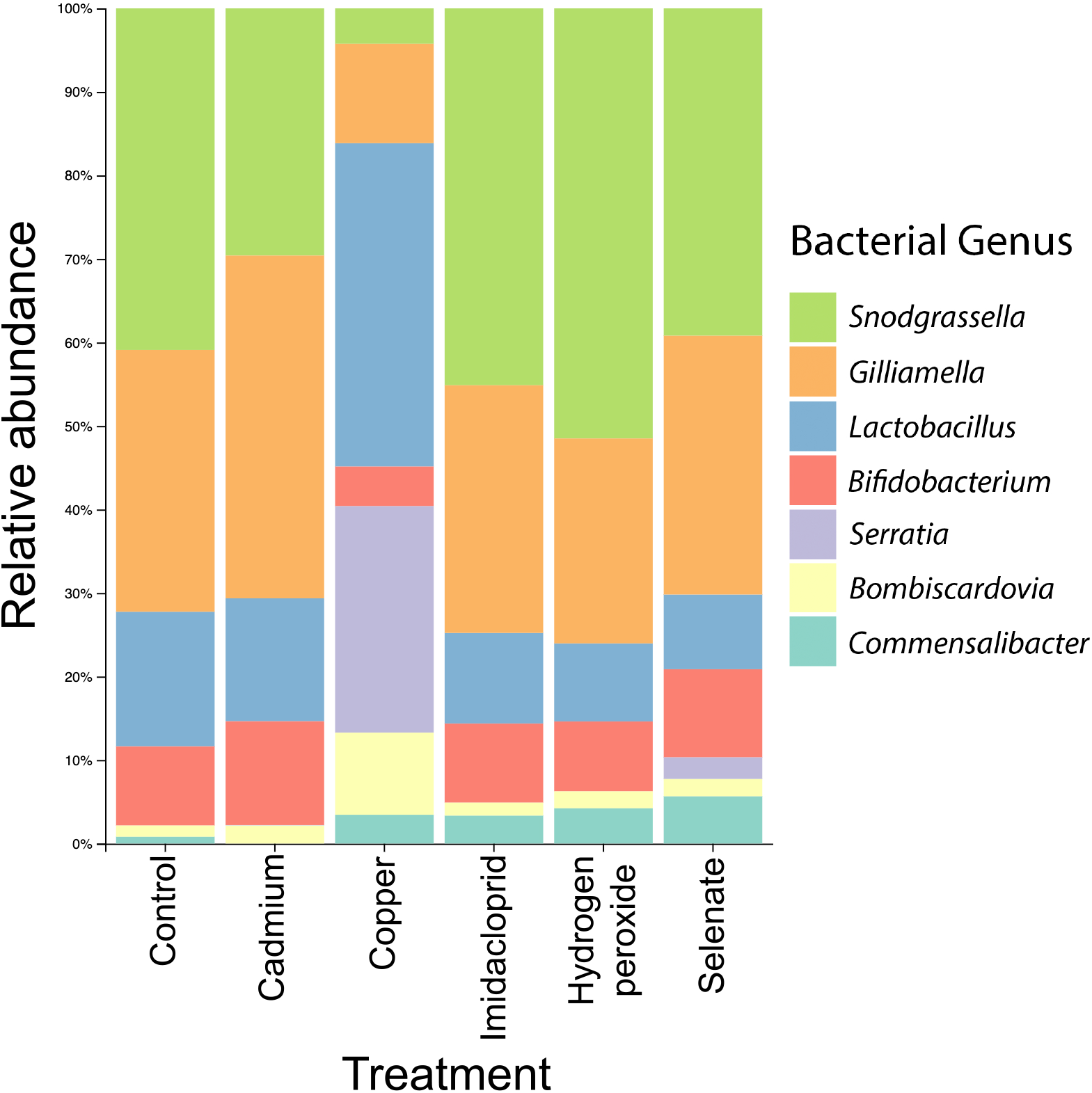
Stacked bar plot showing the relative abundance of bacterial genera present in bumblebee microbiomes following four days of exposure to the various chemicals. Rare genera (<1% relative abundance) were removed for clarity.

**Figure 4.**
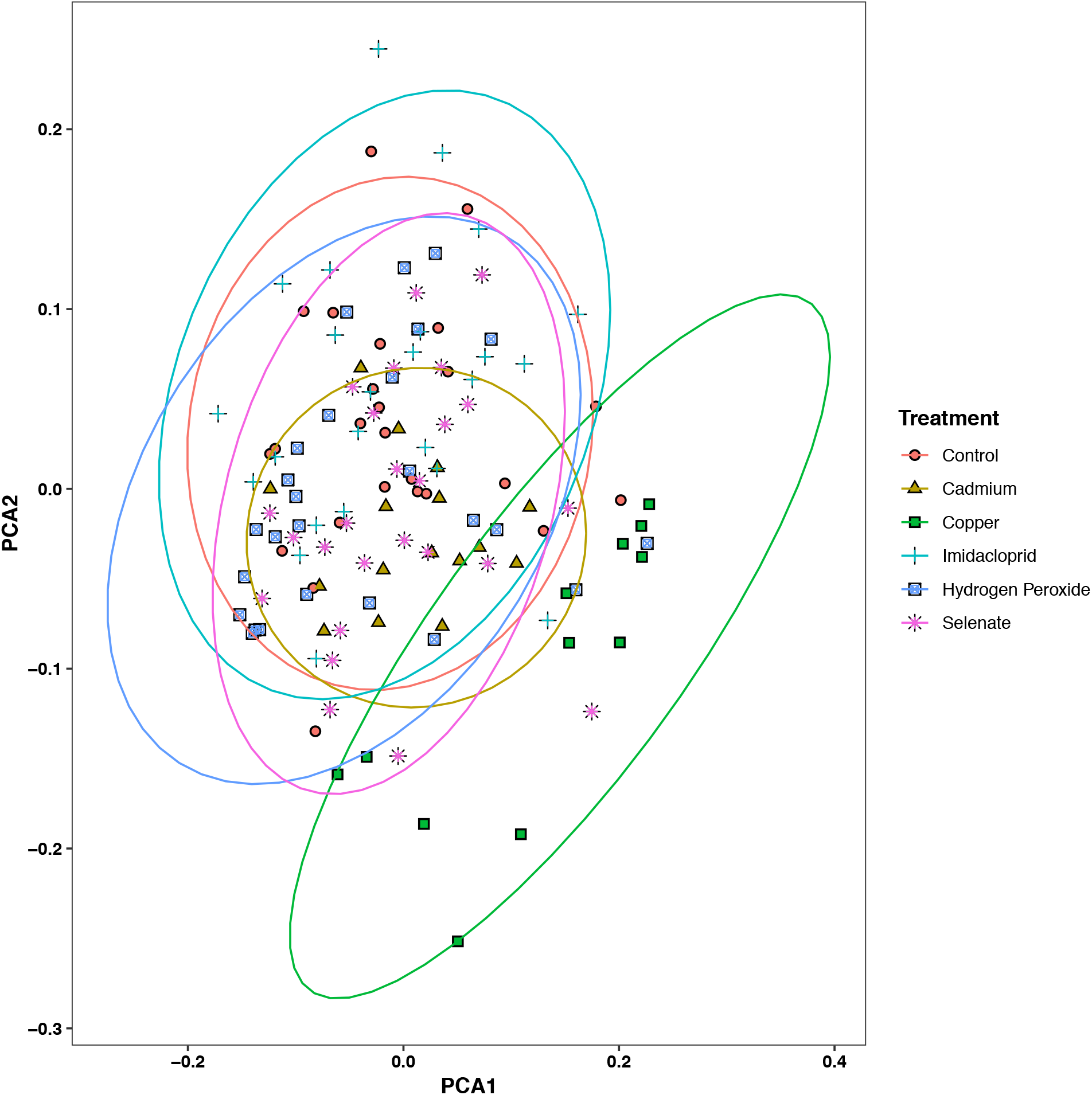
PCoA plot of the Generalized UniFrac distance matrix of bumble bee microbiome samples following a four day chemical exposure by the bee. Overall, treatment (F = 4.57, R^2^ = 0.14, P < 0.001), colony (F = 6.71, R^2^ = 0.08, P < 0.001), and an interaction of these factors (F = 1.63, R^2^ = 0.10, P < 0.001) significantly affected the microbiomes of our samples. Post-hoc testing showed that each treatment except imidacloprid significantly altered the beta diversity of the bees’ microbiomes (BH corrected P_adj_ < 0.02; imidacloprid: P_adj_ = 0.96). Ellipses represent 95% confidence intervals around the centroid of microbiomes for each treatment and color/shape corresponds to treatment.

We used the R package “DESeq2” to identify differences in proportional abundances of ESVs in our treatments versus controls. Several ESVs significantly differed in proportional abundance versus controls (P_adj_ = < 0.05, Fig. 5, Supplemental Table ST4): In cadmium treatments: one *Commensalibacter* ESV was lower; copper treatments: two *Serratia* ESVs were higher, four *Gilliamella* ESVs (two higher and two lower), two *Bombiscardovia* ESVs were higher, one *Commensalibacter* ESV was higher, two *Lactobacillus* ESVs were higher, and two *Snodgrassella* ESVs were lower; hydrogen peroxide: one *Commensalibacter* ESV was higher; selenate treatments: two *Commensalibacter* ESVs were higher, and two *Lactobacillus*, two *Snodgrassella*, and two *Gilliamella* ESVs were all lower; lastly, we did not find any differentially abundant ESVs in our imidacloprid-treated bees.

**Figure 5.**
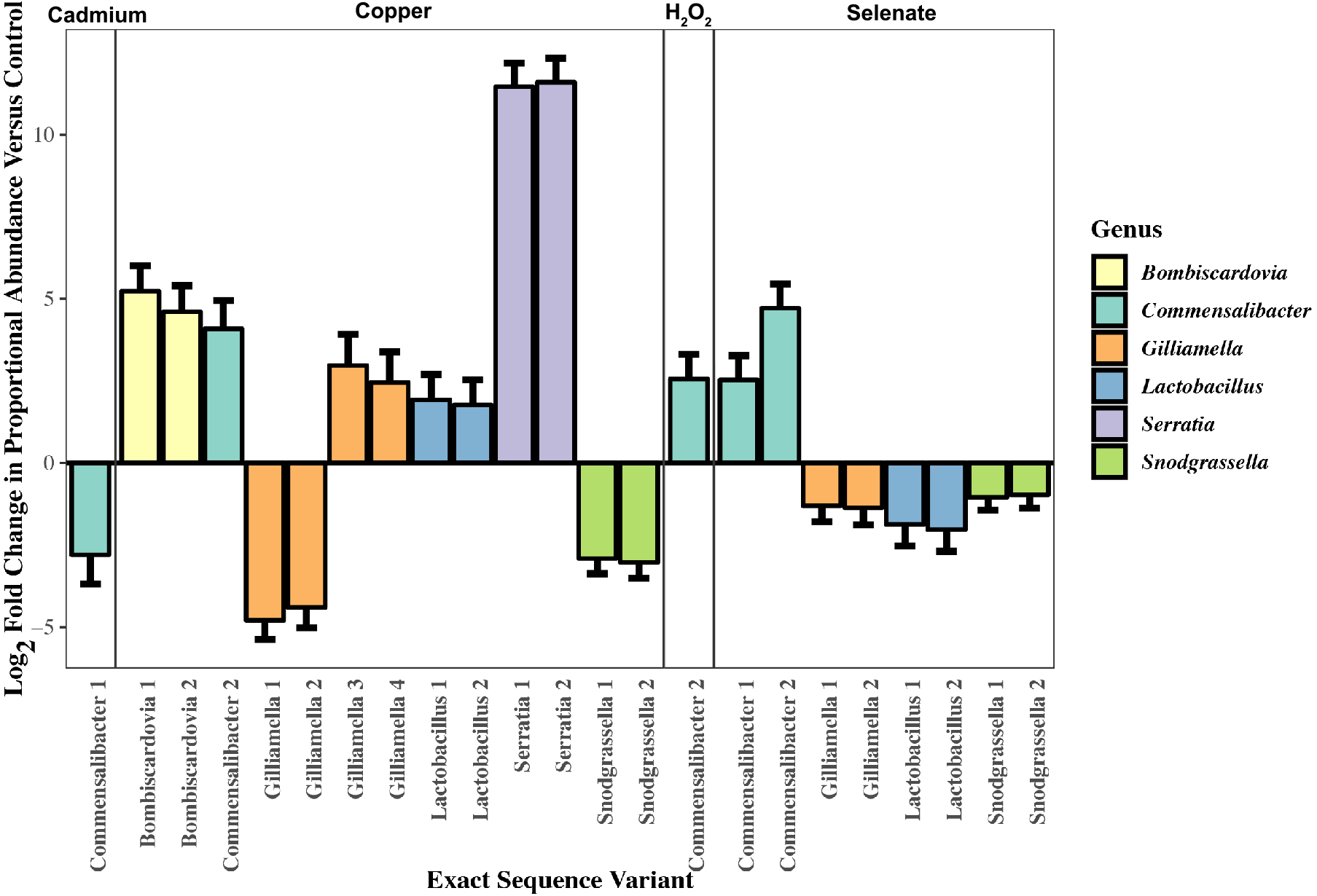
Log_2_fold change of the differentially abundant Exact Sequence Variants (ESV) in the bumble bee microbiomes following exposure to either cadmium, copper, hydrogen peroxide, or selenate treatments versus controls, colored by genus. Each treatment had at least one significantly different ESV except imidacloprid (BH corrected P_adj_ < 0.05). Error bars denote the standard error of the Log_2_fold change.

## Discussion

A range of environmental toxicants negatively influenced bee health indirectly and directly by perturbing their microbiome composition and reducing their survival respectively. Field realistic doses of cadmium, selenate, and copper impacted the bumble bee microbiome - potentially having an indirect negative effect on bumblebee health. Furthermore, there are individual ESVs of symbiotic or pathogenic bacteria that are tolerant or susceptible to most of these chemicals. Previous studies have examined whether the microbiome is affected by our assayed poisons in several non-bee species (42–47) and in honey bees and bumble bees (22,48,53). We extend this work by screening a broad panel of toxicants on bees and their symbionts, and further show that members of the bee microbiota vary in their tolerance to the chemicals. Additionally, we identify that the field realistic concentrations of cadmium and selenate can cause mortality in the common eastern bumble bee, *Bombus impatiens*.

### Toxicants generally, but sometimes subtly, affect the bumble bee microbiome

The bumble bee gut microbiome exhibited a variety of responses to the toxicant challenges. Copper led to a striking proportional increase of the opportunistic pathogen *Serratia*, which suggests a departure from the normal and presumably healthy gut community, potentially resulting in gut dysbiosis (35,98). Selenate exposure allowed non-core bacteria to proliferate, while core symbiont ESVs were less proportionally abundant, further supporting our hypothesis of dysbiosis resulting from toxicant exposure (99). Despite less extreme effects on survival, copper exposure had the most dramatic effect on the bees’ overall microbial diversity and changed the proportional abundance of 13 individual ESVs: most taxa increased in abundance. Conversely, two *G. apicola* ESVs and two *S. alvi* ESVs decreased in abundance. The effect on *G. apicola* is especially interesting, as two other *G. apicola* ESVs significantly increased in abundance, suggesting there is genomic variation in copper tolerance within this taxon, similar to other genomic differences within bee symbionts (96). As a caveat, due to the compositional nature of amplicon-based microbiome sequencing (100), we may be observing changes in the overall microbial population and the treatments may not be causing dramatic effects on individual ESVs.

Genomic analyses suggest putative mechanisms by which the bumble bee gut microbiome is affected by copper and selenate. Each core symbiont varies in its complement of selenium ion resistance genes, with *Bifidobacterium bombi*, *Bombiscardovia coagulans*, *L. bombicola*, and *S. alvi* (97,101,102) all possessing the selenate transporter DedA (75), while *G. apicola* does not. All annotated strains of *S. alvi* possess the sulfate/selenium ion transporter CysA, while some strains of *G. apicola* possess this gene, along with the selenium ion transporter TsgA (76). There is variation in selenium ion transport genes between strains of *G. apicola*, with strains having zero or one copy of CysA, and zero to three copies of TsgA. Similar to selenium tolerance genes, between-strain variation exists in copper tolerance in *S. alvi* and *G. apicola*: Strains of *S. alvi* contain varying numbers of the genes CueO (83), CueR (86), and a copper-translocating ATPase, while *G. apicola* strains varied in SufI and copper-translocating ATPase genes. The strain variation in *G. apicola* copper translocating genes may underlie the differential abundance of *G. apicola* strains under copper challenge, although our 16S rRNA gene data do not allow us to test this hypothesis.

Cadmium, imidacloprid, and hydrogen peroxide all had moderate (cadmium and hydrogen peroxide) to no (imidacloprid) effects on the microbiome. Cadmium changed the bumble bees’ bacterial community but resulted in decreased proportional abundance of only one ESV of *Commensalibacter.* Imidacloprid did not affect the diversity of the gut microbiome in *B. impatiens* whatsoever. These results agree with a previous experiment that showed imidacloprid did not affect the honey bee microbial community, and that bee gut bacteria do not appear to metabolize the neurotoxicant (48). As imidacloprid targets acetylcholine receptors in insects (103) it is perhaps not surprising that the bumble bee gut microbiome is not affected by this insecticide. Hydrogen peroxide modestly changed the microbial community of *B. impatiens* at higher-than-natural concentrations and increased the proportional abundance of one ESV of *Commensalibacter*. As hydrogen peroxide is thought to have antimicrobial properties in flower nectar (104), bumble bee-associated microbes may be resistant due to routine peroxide exposure. The ubiquity of hydrogen peroxide exposure in nature may explain why members of the core bee gut microbiome have various combinations of genes to cope with oxidative stress. While *S. alvi* did not exhibit any genetic strain variation in oxidative stress response, *G. apicola* did: our genomic analysis indicated a variable presence of SoxS, an Fnr regulator, and NnrS. Cadmium resistance is less clear, as *Commensalibacter intestini* has several cadmium resistance genes, but is still susceptible to the treatment *in vivo,* while core bumble bee symbionts’ resistance pathways are more depauperate. As with other resistance pathways, *G. apicola* had varied in cadmium tolerance, while *S. alvi* exhibited substantial strain variation, notably in the cadmium-responsive regulator CadR. These results suggest that individual core bee microbiome members largely resist cadmium on a community-level scale, and we hypothesize that bacteria may be partitioning cadmium detoxification between each other, as has been shown in other metabolic processes (97).

### Mortality effects of each compound

By exposing bees to cadmium, copper, hydrogen peroxide, imidacloprid or selenate, we show that each toxicant is lethal to bumble bees at varying concentrations - following the mantra that the dose makes the poison. For example, constant ingestion of selenate and cadmium at levels that bees may encounter on flowers grown in polluted soils are toxic even on the third day of chronic exposure (105,106). Bees were more tolerant of copper, with lethal doses higher than the levels likely encountered when foraging on plants in contaminated areas (106). In regard to non-elemental toxicants, the insecticide imidacloprid and hydrogen peroxide were both lethal to bees at doses above normal exposure, and we note that bees appeared to avoid the highest doses of hydrogen peroxide. While adult bees tolerated above-field-relevant doses of copper and imidacloprid, sublethal exposure to these chemicals is known to reduce brood production and larvae population, which may cause negative colony-level effects (21,107). Lastly, bees seemed to tolerate natural levels of hydrogen peroxide, which is supported by studies showing high hydrogen peroxide levels in some flowers (16) and that bees can detoxify peroxide (108,109). Our data suggest that exposure to these chemicals should be investigated further, and studies should focus on interactions between bees, gut microbes, parasites, and their environment, to understand more about the subtle and potentially synergistic effects of stressors on pollinator health.

## Conclusion

Bees have been recognized for their use as bioindicators to monitor environmental pollution (110), and our work supports this claim by showing that bees are susceptible to many environmental toxicants. Our interdisciplinary study reports the direct effects of cadmium, copper, hydrogen peroxide, imidacloprid, and selenate exposure, and we conclude that direct effects are only part of the story. To fully appreciate the risks of exposure we must also consider the effects on the microbiome as indirect effects on bee health. Encouragingly, we have identified several potential genomic bases for microbial tolerance or susceptibility to each toxicant and found that there can be substantial between-strain variation in these genes in the core gut bacteria *S. alvi* and *G. apicola*. This variation suggests that the bee gut microbiome harbors diverse strains that may be resilient to various environmental challenges. As we have indicated, there is a wide diversity of stress response genes between bee symbiont strains, and culture-based toxicology assays should be conducted to characterize their individual susceptibility to toxicants *in vitro*. We suggest that future studies investigate the multipartite interactions between host, symbiont, and the environment, and the potential for microbiomes and hosts to reciprocally protect each other from environmental insults.

## Acknowledgements

The authors would like to thank the UC Riverside Genomics Core facility staff for their Next-Generation Sequencing expertise and to Dr. Richard Gill and group for constructive feedback on the manuscript.

## References

1. Garibaldi LA, Steffan-Dewenter I, Winfree R, Aizen MA, Bommarco R, Cunningham SA, et al. Wild pollinators enhance fruit set of crops regardless of honey bee abundance. Science (80-). 2013 Mar 29;340(6127):1608–11. Available from: 10.1126/science.1230200

2. Velthuis HHW. The historical background of the domestication of the bumble-bee, Bombus terrestris, and its introduction in agriculture. Pollination Bees - The Conservation Link Between Agriculture and Nature. 2002.

3. Goulson D, Lye GC, Darvill B. Decline and conservation of bumble bees. Annu Rev Entomol. 2008 Jan 7;53(1):191–208. Available from: 10.1146/annurev.ento.53.103106.093454

4. Colla SR, Gadallah F, Richardson LL, Wagner D, Gall L. Assessing declines of North American bumble bees (*Bombus* spp.) using museum specimens. Biodivers Conserv. 2012 Dec 11;21(14):3585–95. Available from: 10.1007/s10531-012-0383-2

5. Goulson D, Nicholls E, Botias C, Rotheray EL. Bee declines driven by combined stress from parasites, pesticides, and lack of flowers. Science (80-). 2015 Mar 27;347(6229):1255957–1255957. Available from: 10.1126/science.1255957

6. Graystock P, Blane EJ, McFrederick QS, Goulson D, Hughes WOH. Do managed bees drive parasite spread and emergence in wild bees? Int J Parasitol Parasites Wildl. 2016;5(1):64–75. Available from: 10.1016/j.ijppaw.2015.10.001

7. Hladun KR, Di N, Liu TX, Trumble JT. Metal contaminant accumulation in the hive: Consequences for whole-colony health and brood production in the honey bee (*Apis mellifera* L.). Environ Toxicol Chem. 2015;35(2):322–9. Available from: 10.1002/etc.3273

8. Park S, Thornburg RW. Biochemistry of nectar proteins. Vol. 52, Journal of Plant Biology. Springer; 2009. p. 27–34. Available from: 10.1007/s12374-008-9007-5

9. Hladun KR, Parker DR, Trumble JT. Selenium accumulation in the floral tissues of two Brassicaceae species and its impact on floral traits and plant performance. Environ Exp Bot. 2011;74(0):90–7. Available from: 10.1016/j.envexpbot.2011.05.003

10. Sovik E, Perry CJ, LaMora A, Barron AB, Ben-Shahar Y. Negative impact of manganese on honeybee foraging. Biol Lett. 2015;11(3). Available from: 10.1098/rsbl.2014.0989

11. Goulson D. An overview of the environmental risks posed by neonicotinoid insecticides. Vol. 50, Journal of Applied Ecology. 2013. p. 977–87. Available from: 10.1111/1365-2664.12111

12. Vickerman DB, Trumble JT, George GN, Pickering IJ, Nichol H. Selenium biotransformations in an insect ecosystem: Effects of insects on phytoremediation. Environ Sci Technol. 2004 Jul;38(13):3581–6. Available from: 10.1021/es049941s

13. Hutchinson TC, Whitby LM. Heavy-metal pollution in the sudbury mining and smelting region of canada, I. soil and vegetation contamination by nickel, copper, and other metals. Environ Conserv. 1974 Jun 24;1(02):123. Available from: 10.1017/S0376892900004240

14. Epstein L, Bassein S. Pesticide applications of copper on perennial crops in California, 1993 to 1998. J Environ Qual. 2001;(30):1844–7.

15. Tchounwou PB, Yedjou CG, Patlolla AK, Sutton DJ. Heavy metal toxicity and the environment. Vol. 101, EXS. NIH Public Access; 2012. 133–164 p. Available from: 10.1007/978-3-7643-8340-4_6

16. Carter C, Thornburg RW. Is the nectar redox cycle a floral defense against microbial attack? Trends Plant Sci. 2004 Jul 1;9(7):320–4. Available from: 10.1016/j.tplants.2004.05.008

17. Boyd RS, Martens SN. The significance of metal hyperaccumulation for biotic interactions. Chemoecology. 1998;8(1):1–7. Available from: 10.1007/s000490050002

18. Conti ME, Botrè F. Honeybees and their products as potential bioindicators of heavy metals contamination. Environ Monit Assess. 2001; Available from: 10.1023/A:1010719107006

19. Heard MS, Baas J, Dorne J Lou, Lahive E, Robinson AG, Rortais A, et al. Comparative toxicity of pesticides and environmental contaminants in bees: Are honey bees a useful proxy for wild bee species? Sci Total Environ. 2017 Feb 1;578:357–65. Available from: 10.1016/j.scitotenv.2016.10.180

20. Hladun KR, Smith BH, Mustard JA, Morton RR, Trumble JT. Selenium toxicity to honey bee (*Apis mellifera* L.) pollinators: effects on behaviors and survival. PLoS One. 2012;7(4):e34137. Available from: 10.1371/journal.pone.0034137

21. Di N, Hladun KR, Zhang K, Liu TX, Trumble JT. Laboratory bioassays on the impact of cadmium, copper and lead on the development and survival of honeybee *(Apis mellifera L.)* larvae and foragers. Chemosphere. 2016;152:530–8. Available from: 10.1016/j.chemosphere.2016.03.033

22. Rothman JA, Leger L, Kirkwood JS, McFrederick QS. Cadmium and selenate exposure affects the honey bee microbiome and metabolome, and bee-associated bacteria show potential for bioaccumulation. Appl Environ Microbiol. 2019 Aug 30;85(21). Available from: 10.1128/AEM.01411-19

23. Engel P, Kwong WK, McFrederick QS, Anderson KE, Barribeau SM, Chandler JA, et al. The bee microbiome: Impact on bee health and model for evolution and ecology of host-microbe interactions. MBio. 2016;7(2):e02164–15. Available from: 10.1128/mBio.02164-15

24. Koch H, Schmid-Hempel P. Bacterial communities in central european bumblebees: low diversity and high specificity. Microb Ecol. 2011 Jul 10;62(1):121–33. Available from: 10.1007/s00248-011-9854-3

25. Koch H, Schmid-Hempel P. Socially transmitted gut microbiota protect bumble bees against an intestinal parasite. Proc Natl Acad Sci USA. 2011;108(48):19288–92. Available from: 10.1073/pnas.1110474108

26. Martinson VG, Danforth BN, Minckley RL, Rueppell O, Tingek S, Moran NA. A simple and distinctive microbiota associated with honey bees and bumble bees. Mol Ecol. 2011;20(3):619–28. Available from: 10.1111/j.1365-294X.2010.04959.x

27. Kwong WK, Medina LA, Koch H, Sing K-W, Soh EJY, Ascher JS, et al. Dynamic microbiome evolution in social bees. Sci Adv. 2017 Mar 30;3(3):e1600513. Available from: 10.1126/sciadv.1600513

28. Powell JE, Ratnayeke N, Moran NA. Strain diversity and host specificity in a specialized gut symbiont of honeybees and bumblebees. Mol Ecol. 2016 Sep;25(18):4461–71. Available from: 10.1111/mec.13787

29. Zheng H, Powell JE, Steele MI, Dietrich C, Moran NA. Honeybee gut microbiota promotes host weight gain via bacterial metabolism and hormonal signaling. Proc Natl Acad Sci USA. 2017 May 2;114(18):4775–80. Available from: 10.1073/pnas.1701819114

30. Kwong WK, Mancenido AL, Moran NA. Immune system stimulation by the native gut microbiota of honey bees. R Soc Open Sci. 2017 Mar 7;4(2):170003. Available from: 10.1098/rsos.170003

31. Palmer-Young EC, Raffel TR, McFrederick QS. Temperature-mediated inhibition of a bumblebee parasite by an intestinal symbiont. Proc R Soc B Biol Sci. 2018 Nov 7;285(1890):20182041. Available from: 10.1098/rspb.2018.2041

32. Schwarz RS, Moran NA, Evans JD. Early gut colonizers shape parasite susceptibility and microbiota composition in honey bee workers. Proc Natl Acad Sci U S A. 2016;113(33):9345–50. Available from: 10.1073/pnas.1606631113

33. Rothman JA, Carroll MJ, Meikle WG, Anderson KE, McFrederick QS. Longitudinal effects of supplemental forage on the honey bee (Apis mellifera) microbiota and inter- and intra-colony variability. Microb Ecol. 2018 Feb 3;76(3):814–24. Available from: 10.1007/s00248-018-1151-y

34. Kešnerová L, Emery O, Troilo M, Liberti J, Erkosar B, Engel P. Gut microbiota structure differs between honeybees in winter and summer. ISME J. 2019 Dec 13; Available from: 10.1038/s41396-019-0568-8

35. Raymann K, Shaffer Z, Moran NA. Antibiotic exposure perturbs the gut microbiota and elevates mortality in honeybees. PLoS Biol. 2017;15(3). Available from: 10.1371/journal.pbio.2001861

36. Rubanov A, Russell KA, Rothman JA, Nieh JC, McFrederick QS. Intensity of *Nosema ceranae* infection is associated with specific honey bee gut bacteria and weakly associated with gut microbiome structure. Sci Rep. 2019 Dec 7;9(1):3820. Available from: 10.1038/s41598-019-40347-6

37. Kakumanu ML, Reeves AM, Anderson TD, Rodrigues RR, Williams MA. Honey bee gut microbiome is altered by in-hive pesticide exposures. Front Microbiol. 2016 Aug 16;7:1255. Available from: 10.3389/fmicb.2016.01255

38. Maes P, Rodrigues P, Oliver R, Mott BM, Anderson KE. Diet related gut bacterial dysbiosis correlates with impaired development, increased mortality and *Nosema* disease in the honey bee *Apis mellifera*. Mol Ecol. 2016;25(21):5439–50. Available from: 10.1111/mec.13862

39. Engel P, Bartlett KD, Moran NA. The bacterium *Frischella perrara* causes scab formation in the gut of its honeybee host. MBio. 2015 May 19;6(3):1–8. Available from: 10.1128/mBio.00193-15

40. Cornman RS, Tarpy DR, Chen Y, Jeffreys L, Lopez D, Pettis JS, et al. Pathogen webs in collapsing honey bee colonies. PLoS One. 2012;7(8):e43562. Available from: 10.1371/journal.pone.0043562

41. Claus SP, Guillou H, Ellero-Simatos S. The gut microbiota: a major player in the toxicity of environmental pollutants? npj Biofilms Microbiomes. 2016 May 10;2(1):16003. Available from: 10.1038/npjbiofilms.2016.3

42. Breton J, Massart S, Vandamme P, De Brandt E, Pot B, Foligné B. Ecotoxicology inside the gut: impact of heavy metals on the mouse microbiome. BMC Pharmacol Toxicol. 2013;14:62. Available from: 10.1186/2050-6511-14-62

43. Chang X, Li H, Feng J, Chen Y, Nie G, Zhang J. Effects of cadmium exposure on the composition and diversity of the intestinal microbial community of common carp (*Cyprinus carpio* L.). Ecotoxicol Environ Saf. 2019 Apr 30;171:92–8. Available from: 10.1016/J.ECOENV.2018.12.066

44. Kasaikina M V, Kravtsova MA, Lee BC, Seravalli J, Peterson DA, Walter J, et al. Dietary selenium affects host selenoproteome expression by influencing the gut microbiota. FASEB J. 2011 Jul 1;25(7):2492–9. Available from: 10.1096/fj.11-181990

45. Zhai Q, Li T, Yu L, Xiao Y, Feng S, Wu J, et al. Effects of subchronic oral toxic metal exposure on the intestinal microbiota of mice. Sci Bull. 2017 Jun 30;62(12):831–40. Available from: 10.1016/J.SCIB.2017.01.031

46. Daisley BA, Trinder M, McDowell TW, Welle H, Dube JS, Ali SN, et al. Neonicotinoid-induced pathogen susceptibility is mitigated by *Lactobacillus plantarum* immune stimulation in a *Drosophila melanogaster* model. Sci Rep. 2017 Dec 2;7(1):2703. Available from: 10.1038/s41598-017-02806-w

47. Oliveira JHM, Gonçalves RLS, Lara FA, Dias FA, Gandara ACP, Menna-Barreto RFS, et al. blood meal-derived heme decreases ROS levels in the midgut of *Aedes aegypti* and allows proliferation of intestinal microbiota. Schneider DS, editor. PLoS Pathog. 2011 Mar 17;7(3):e1001320. Available from: 10.1371/journal.ppat.1001320

48. Raymann K, Motta EVS, Girard C, Riddington IM, Dinser JA, Moran NA. Imidacloprid decreases honey bee survival rates but does not affect the gut microbiome. Appl Environ Microbiol. 2018 Apr 20;84(13):AEM.00545–18. Available from: 10.1128/AEM.00545-18

49. Senderovich Y, Halpern M. The protective role of endogenous bacterial communities in chironomid egg masses and larvae. ISME J. 2013;7(11):2147–58. Available from: 10.1038/ismej.2013.100

50. Tian F, Xiao Y, Li X, Zhai Q, Wang G, Zhang Q, et al. protective effects of *Lactobacillus plantarum* CCFM8246 against copper toxicity in mice. Bach H, editor. PLoS One. 2015 Nov 25;10(11):e0143318. Available from: 10.1371/journal.pone.0143318

51. Coryell M, McAlpine M, Pinkham N V., McDermott TR, Walk ST. The gut microbiome is required for full protection against acute arsenic toxicity in mouse models. Nat Commun. 2018 Dec 21;9(1):5424. Available from: 10.1038/s41467-018-07803-9

52. Wang Y, Shu X, Zhou Q, Fan T, Wang T, Chen X, et al. Selenite reduction and the biogenesis of selenium nanoparticles by *Alcaligenes faecalis* se03 isolated from the gut of *Monochamus alternatus* (Coleoptera: Cerambycidae). Int J Mol Sci. 2018;19(9). Available from: 10.3390/ijms19092799

53. Rothman JA, Leger L, Graystock P, Russell K, McFrederick QS. The bumble bee microbiome increases survival of bees exposed to selenate toxicity. Environ Microbiol. 2019 Sep 14;21(9):3417–29. Available from: 10.1111/1462-2920.14641

54. Aziz RK, Bartels D, Best AA, DeJongh M, Disz T, Edwards RA, et al. The RAST Server: rapid annotations using subsystems technology. BMC Genomics. 2008;9:75. Available from: 10.1186/1471-2164-9-75

55. Christian R, Strebig JC. Package “drc”: Analysis of Dose-Response Curves. 2016.

56. Therneau TM. A Package for Survival Analysis in S. 2015.

57. Kassambara A, Kosinski M. survminer: Drawing Survival Curves using “ggplot2.” 2018.

58. Engel P, James RR, Koga R, Kwong WK, McFrederick QS, Moran NA. Standard methods for research on *Apis mellifera* gut symbionts. J Apic Res. 2013 Jan 2;52(4):1–24. Available from: 10.3896/IBRA.1.52.4.07

59. Pennington MJ, Rothman JA, Jones MB, McFrederick QS, Gan J, Trumble JT. Effects of contaminants of emerging concern on *Megaselia scalaris* (Lowe, Diptera: Phoridae) and its microbial community. Sci Rep. 2017 Dec 15;7(1):8165. Available from: 10.1038/s41598-017-08683-7

60. Rothman JA, Andrikopoulos C, Cox-Foster D, McFrederick QS. Floral and foliar source affect the bee nest microbial community. Microb Ecol. 2019 Dec 14;78(2):506–16. Available from: 10.1007/s00248-018-1300-3

61. McFrederick QS, Rehan SM. Characterization of pollen and bacterial community composition in brood provisions of a small carpenter bee. Mol Ecol. 2016;25(10):2302–11. Available from: 10.1111/mec.13608

62. Pennington MJ, Rothman JA, Jones MB, McFrederick QS, Gan J, Trumble JT. Effects of contaminants of emerging concern on *Myzus persicae* (Sulzer, Hemiptera: Aphididae) biology and on their host plant, *Capsicum annuum*. Environ Monit Assess. 2018 Mar 8;190(3):125. Available from: 10.1007/s10661-018-6503-z

63. Kembel SW, O’Connor TK, Arnold HK, Hubbell SP, Wright SJ, Green JL. Relationships between phyllosphere bacterial communities and plant functional traits in a neotropical forest. Proc Natl Acad Sci USA. 2014;111(38):13715–20. Available from: 10.1073/pnas.1216057111

64. Hanshew AS, Mason CJ, Raffa KF, Currie CR. Minimization of chloroplast contamination in 16S rRNA gene pyrosequencing of insect herbivore bacterial communities. J Microbiol Methods. 2013;95(2):149–55. Available from: 10.1016/j.mimet.2013.08.007

65. Bolyen E, Rideout JR, Dillon MR, Bokulich NA, Abnet CC, Al-Ghalith GA, et al. Reproducible, interactive, scalable and extensible microbiome data science using QIIME 2. Nat Biotechnol. 2019 Jul 24; Available from: 10.1038/s41587-019-0209-9

66. Callahan BJ, McMurdie PJ, Rosen MJ, Han AW, Johnson AJA, Holmes SP. DADA2: High-resolution sample inference from Illumina amplicon data. Nat Methods. 2016 Jul 23;13(7):581–3. Available from: 10.1038/nmeth.3869

67. Bokulich NA, Kaehler BD, Rideout JR, Dillon M, Bolyen E, Knight R, et al. Optimizing taxonomic classification of marker-gene amplicon sequences with QIIME 2’s q2-feature-classifier plugin. Microbiome. 2018 Dec 17;6(1):90. Available from: 10.1186/s40168-018-0470-z

68. Quast C, Pruesse E, Yilmaz P, Gerken J, Schweer T, Yarza P, et al. The SILVA ribosomal RNA gene database project: Improved data processing and web-based tools. Nucleic Acids Res. 2013 Nov 27;41(D1):D590–6. Available from: 10.1093/nar/gks1219

69. Katoh K, Standley DM. MAFFT multiple sequence alignment software version 7: improvements in performance and usability. Mol Biol Evol. 2013 Apr;30(4):772–80. Available from: 10.1093/molbev/mst010

70. Price MN, Dehal PS, Arkin AP. FastTree 2--approximately maximum-likelihood trees for large alignments. PLoS One. 2010;5(3):e9490.

71. Wickham H. ggplot2: Elegant graphics for data analysis. Springer-Verlag New York; 2009.

72. R Core Team. R: A language and environment for statistical computing. Vienna, Austria; 2018.

73. Oksanen J, Blanchet FG, Friendly M, Kindt R, Legendre P, McGlinn D, et al. vegan: Community Ecology Package. 2017.

74. Love MI, Huber W, Anders S. Moderated estimation of fold change and dispersion for RNA-seq data with DESeq2. Genome Biol. 2014 Dec 5;15(12):550. Available from: 10.1186/s13059-014-0550-8

75. Ledgham F, Quest B, Vallaeys T, Mergeay M, Covès J. A probable link between the DedA protein and resistance to selenite. Res Microbiol. 2005 Sep 2;156(3):367–74. Available from: 10.1016/j.resmic.2004.11.003

76. Guzzo J, Dubow MS. A novel selenite- and tellurite-inducible gene in *Escherichia coli*. Appl Environ Microbiol. 2000 Nov 1;66(11):4972–8. Available from: 10.1128/AEM.66.11.4972-4978.2000

77. Lindblow-Kull C, Kull FJ, Shrift A. Single transporter for sulfate, selenate, and selenite in *Escherichia coli* K-12. J Bacteriol. 1985 Sep;163(3):1267–9.

78. Nies DH. The cobalt, zinc, and cadmium efflux system CzcABC from *Alcaligenes eutrophus* functions as a cation-proton antiporter in *Escherichia coli*. J Bacteriol. 1995;177(10):2707–12.

79. Anton A, Grosse C, Reissmann J, Pribyl T, Nies DH. CzcD is a heavy metal ion transporter involved in regulation of heavy metal resistance in *Ralstonia* sp. strain CH34. J Bacteriol. 1999 Nov 15;181(22):6876–81.

80. Brocklehurst K., Megit S., Morby A. Characterisation of CadR from *Pseudomonas aeruginosa*: a Cd(II)-responsive MerR homologue. Biochem Biophys Res Commun. 2003 Aug 22;308(2):234–9. Available from: 10.1016/S0006-291X(03)01366-4

81. Axelsen KB, Palmgren MG. Evolution of substrate specificities in the P-Type ATPase superfamily. J Mol Evol. 1998 Jan;46(1):84–101. Available from: 10.1007/PL00006286

82. Nakamura K, Go N. Function and molecular evolution of multicopper blue proteins. Cell Mol Life Sci. 2005 Sep 9;62(18):2050–66. Available from: 10.1007/s00018-004-5076-x

83. Grass G, Rensing C. Genes Involved in copper homeostasis in *Escherichia coli*. J Bacteriol. 2001 Mar 15;183(6):2145–7. Available from: 10.1128/JB.183.6.2145-2147.2001

84. Gupta SD, Wu HC, Rick PD. A *Salmonella typhimurium* genetic locus which confers copper tolerance on copper-sensitive mutants of *Escherichia coli*. J Bacteriol. 1997;179(16):4977–84. Available from: 10.1128/jb.179.16.4977-4984.1997

85. Hu Y, Wang H, Zhang M, Sun L. Molecular analysis of the copper-responsive CopRSCD of a pathogenic *Pseudomonas fluorescens* strain. J Microbiol. 2009 Jun 26;47(3):277–86. Available from: 10.1007/s12275-008-0278-9

86. Stoyanov J V., Hobman JL, Brown NL. CueR (YbbI) of *Escherichia coli* is a MerR family regulator controlling expression of the copper exporter CopA. Mol Microbiol. 2001 Jan 1;39(2):502–12. Available from: 10.1046/j.1365-2958.2001.02264.x

87. Yesilkaya H, Kadioglu A, Gingles N, Alexander JE, Mitchell TJ, Andrew PW. Role of manganese-containing superoxide dismutase in oxidative stress and virulence of *Streptococcus pneumoniae*. Infect Immun. 2000 May;68(5):2819–26.

88. Wu J, Weiss B. Two divergently transcribed genes, soxR and soxS, control a superoxide response regulon of *Escherichia coli*. J Bacteriol. 1991 May 1;173(9):2864–71. Available from: 10.1128/JB.173.9.2864-2871.1991

89. Maddocks SE, Oyston PCF. Structure and function of the LysR-type transcriptional regulator (LTTR) family proteins. Microbiology. 2008 Dec 1;154(12):3609–23. Available from: 10.1099/mic.0.2008/022772-0

90. Troxell B, Hassan HM. Transcriptional regulation by ferric uptake regulator (Fur) in pathogenic bacteria. Front Cell Infect Microbiol. 2013;3:59. Available from: 10.3389/fcimb.2013.00059

91. Gaballa A, Helmann JD. A peroxide-induced zinc uptake system plays an important role in protection against oxidative stress in *Bacillus subtilis*. Mol Microbiol. 2002 Aug 1;45(4):997–1005. Available from: 10.1046/j.1365-2958.2002.03068.x

92. Stern AM, Liu B, Bakken LR, Shapleigh JP, Zhu J. A novel protein protects bacterial iron-dependent metabolism from nitric oxide. J Bacteriol. 2013 Oct;195(20):4702–8. Available from: 10.1128/JB.00836-13

93. Scott C, Rawsthorne H, Upadhyay M, Shearman CA, Gasson MJ, Guest JR, et al. Zinc uptake, oxidative stress and the FNR-like proteins of *Lactococcus lactis*. FEMS Microbiol Lett. 2000 Nov 1;192(1):85–9. Available from: 10.1111/j.1574-6968.2000.tb09363.x

94. Chelikani P, Fita I, Loewen PC. Diversity of structures and properties among catalases. Cell Mol Life Sci. 2004 Jan 1;61(2):192–208. Available from: 10.1007/s00018-003-3206-5

95. Wang H-W, Chung C-H, Ma T-Y, Wong H. Roles of alkyl hydroperoxide reductase subunit C (AhpC) in viable but nonculturable *Vibrio parahaemolyticus*. Appl Environ Microbiol. 2013 Jun;79(12):3734–43. Available from: 10.1128/AEM.00560-13

96. Engel P, Stepanauskas R, Moran NA. Hidden diversity in honey bee gut symbionts detected by single-cell genomics. PLoS Genet. 2014;10(9):e1004596. Available from: 10.1371/journal.pgen.1004596

97. Kwong WK, Engel P, Koch H, Moran NA. Genomics and host specialization of honey bee and bumble bee gut symbionts. Proc Natl Acad Sci USA. 2014;111(31):11509–14. Available from: 10.1073/pnas.1405838111

98. Raymann K, Coon KL, Shaffer Z, Salisbury S, Moran NA. Pathogenicity of *Serratia marcescens* strains in honey bees. MBio. 2018 Oct 9;9(5):e01649–18. Available from: 10.1128/mBio.01649-18

99. Anderson KE, Ricigliano VA. Honey bee gut dysbiosis: a novel context of disease ecology. Curr Opin Insect Sci. 2017 Aug;22:125–32. Available from: 10.1016/j.cois.2017.05.020

100. Gloor GB, Macklaim JM, Pawlowsky-Glahn V, Egozcue JJ. Microbiome datasets are compositional: and this is not optional. Front Microbiol. 2017 Nov 15;8:2224. Available from: 10.3389/fmicb.2017.02224

101. Martinson VG, Magoc T, Koch H, Salzberg SL, Moran NA. Genomic features of a bumble bee symbiont reflect its host environment. Appl Environ Microbiol. 2014 Jul 1;80(13):3793–803. Available from: 10.1128/AEM.00322-14

102. Milani C, Lugli GA, Duranti S, Turroni F, Bottacini F, Mangifesta M, et al. Genomic encyclopedia of type strains of the genus *Bifidobacterium*. Appl Environ Microbiol. 2014 Oct 15;80(20):6290–302. Available from: 10.1128/AEM.02308-14

103. Matsuda K, Buckingham SD, Kleier D, Rauh JJ, Grauso M, Sattelle DB. Neonicotinoids: insecticides acting on insect nicotinic acetylcholine receptors. Trends Pharmacol Sci. 2001 Nov 1;22(11):573–80. Available from: 10.1016/S0165-6147(00)01820-4

104. González-Teuber M, Heil M. Nectar chemistry is tailored for both attraction of mutualists and protection from exploiters. Plant Signal Behav. 2009 Sep 28;4(9):809–13. Available from: 10.4161/psb.4.9.9393

105. Hladun KR, Parker DR, Tran KD, Trumble JT. Effects of selenium accumulation on phytotoxicity, herbivory, and pollination ecology in radish (*Raphanus sativus* L.). Environ Pollut. 2012 Jan;172:70–5. Available from: 10.1016/j.envpol.2012.08.009

106. Hladun KR, Parker DR, Trumble JT. Cadmium, copper, and lead accumulation and bioconcentration in the vegetative and reproductive organs of *Raphanus sativus*: implications for plant performance and pollination. J Chem Ecol. 2015 Apr 7;41(4):386–95. Available from: 10.1007/s10886-015-0569-7

107. Tasei JN, Lerin J, Ripault G. Sub-lethal effects of imidacloprid on bumblebees, *Bombus terrestris* (Hymenoptera: Apidae), during a laboratory feeding test. Pest Manag Sci. 2000;56(9):784–8. Available from: 10.1002/1526-4998(200009)56:9<784::AID-PS208>3.0.CO;2-T

108. Weirich GF, Collins AM, Williams VP. Antioxidant enzymes in the honey bee, *Apis mellifera*. Apidologie. 2002 Jan;33(1):3–14. Available from: 10.1051/apido:2001001

109. Hu Z, Lee KS, Choo YM, Yoon HJ, Lee SM, Lee JH, et al. Molecular cloning and characterization of 1-Cys and 2-Cys peroxiredoxins from the bumblebee *Bombus ignitus*. Comp Biochem Physiol Part B Biochem Mol Biol. 2010 Mar 1;155(3):272–80. Available from: 10.1016/J.CBPB.2009.11.011

110. Kevan PG. Pollinators as bioindicators of the state of the environment: species, activity and diversity. Vol. 74, Ecosystems and Environment. 1999.

