## Supplemental Information for "The direct and indirect effects of environmental toxicants on the health of bumble bees and their microbiomes"

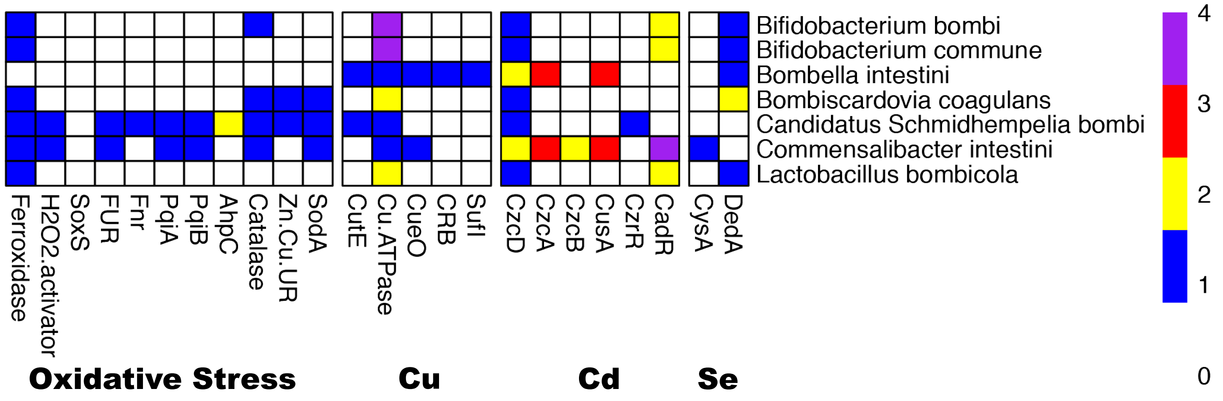


Supplemental Figure S1: Heatmap representing the oxidative stress, copper, cadmium, or selenium ion tolerance genes found in genomes as annotated by RAST. Color represents the copy number of each gene, row names indicate the bacterial species, and column names denote the abbreviated gene name.

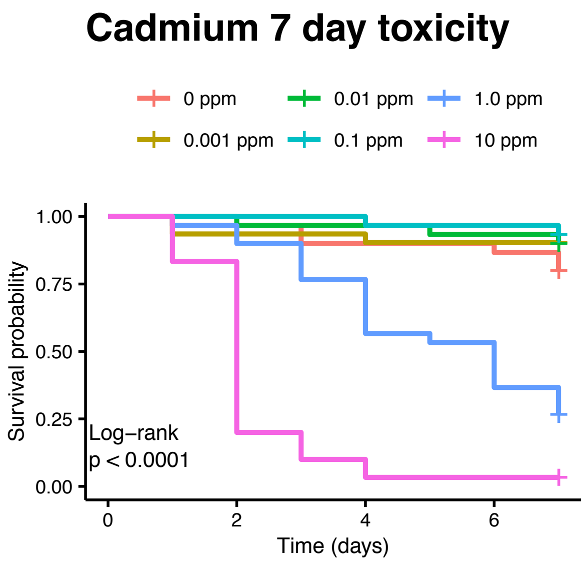
Cadmium 7 day toxicity
0 ppm
0.001 ppm
0.01 ppm
0.1 ppm
1.0 ppm
10 ppm
Survival probability
1.00
0.75
0.50
0.25
0.00
Log–rank
p < 0.0001
0
2
4
6
Time (days)

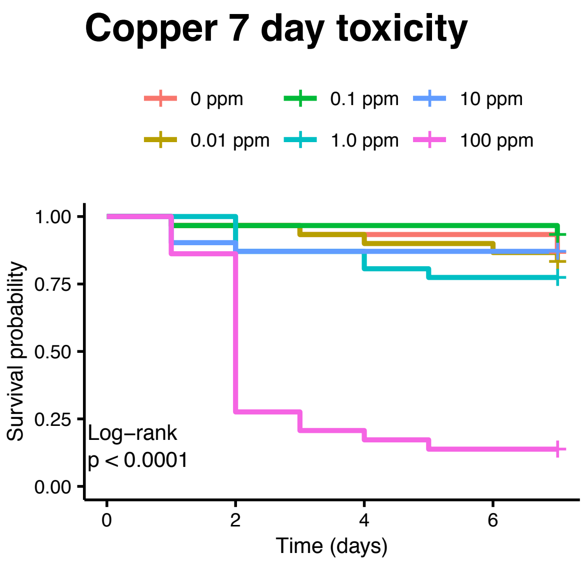
Copper 7 day toxicity
0 ppm
0.01 ppm
0.1 ppm
1.0 ppm
10 ppm
100 ppm
Survival probability
1.00
0.75
0.50
0.25
0.00
Log–rank
p < 0.0001
0
2
4
6
Time (days)

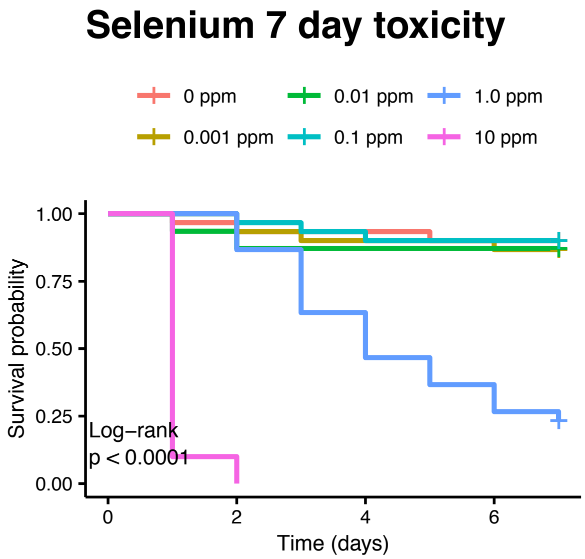
Selenium 7 day toxicity
0 ppm
0.001 ppm
0.01 ppm
0.1 ppm
1.0 ppm
10 ppm
Survival probability
1.00
0.75
0.50
0.25
0.00
Log–rank
p < 0.0001
0
2
4
6
Time (days)

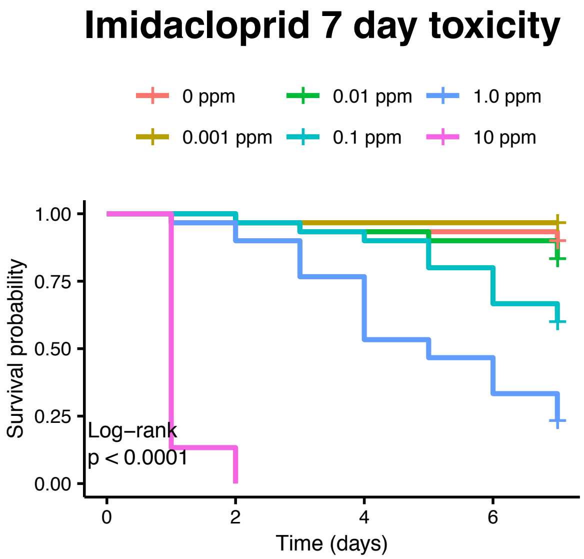
Imidacloprid 7 day toxicity
0 ppm
0.001 ppm
0.01 ppm
0.1 ppm
1.0 ppm
10 ppm
Survival probability
1.00
0.75
0.50
0.25
0.00
Log–rank
p < 0.0001
0
2
4
6
Time (days)

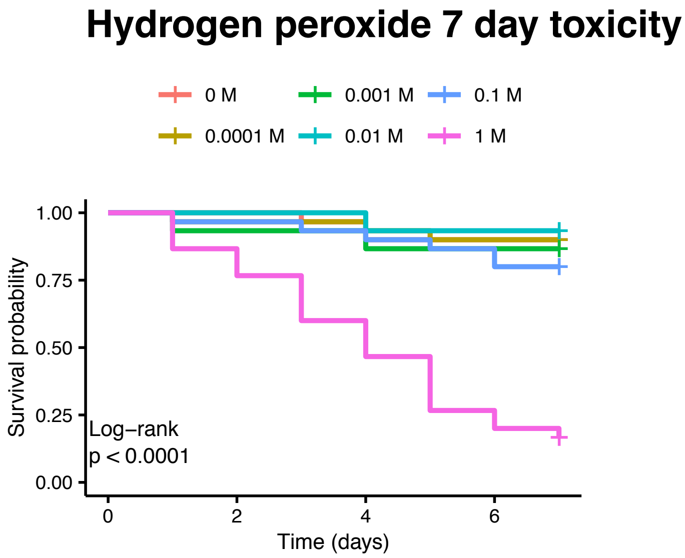
Hydrogen peroxide 7 day toxicity
0 M
0.0001 M
0.001 M
0.01 M
0.1 M
1 M
Survival probability
1.00
0.75
0.50
0.25
0.00
Log–rank
p < 0.0001
0
2
4
6
Time (days)

Supplemental Figure S2: Survival probabilities of *Bombus impatiens* exposed to cadmium, copper, selenate, imidacloprid, and hydrogen peroxide for seven days at each treatment dose. Toxicant exposure significantly reduced bee survivorship (cox proportional hazard test $P < 0.001$ for each compound).

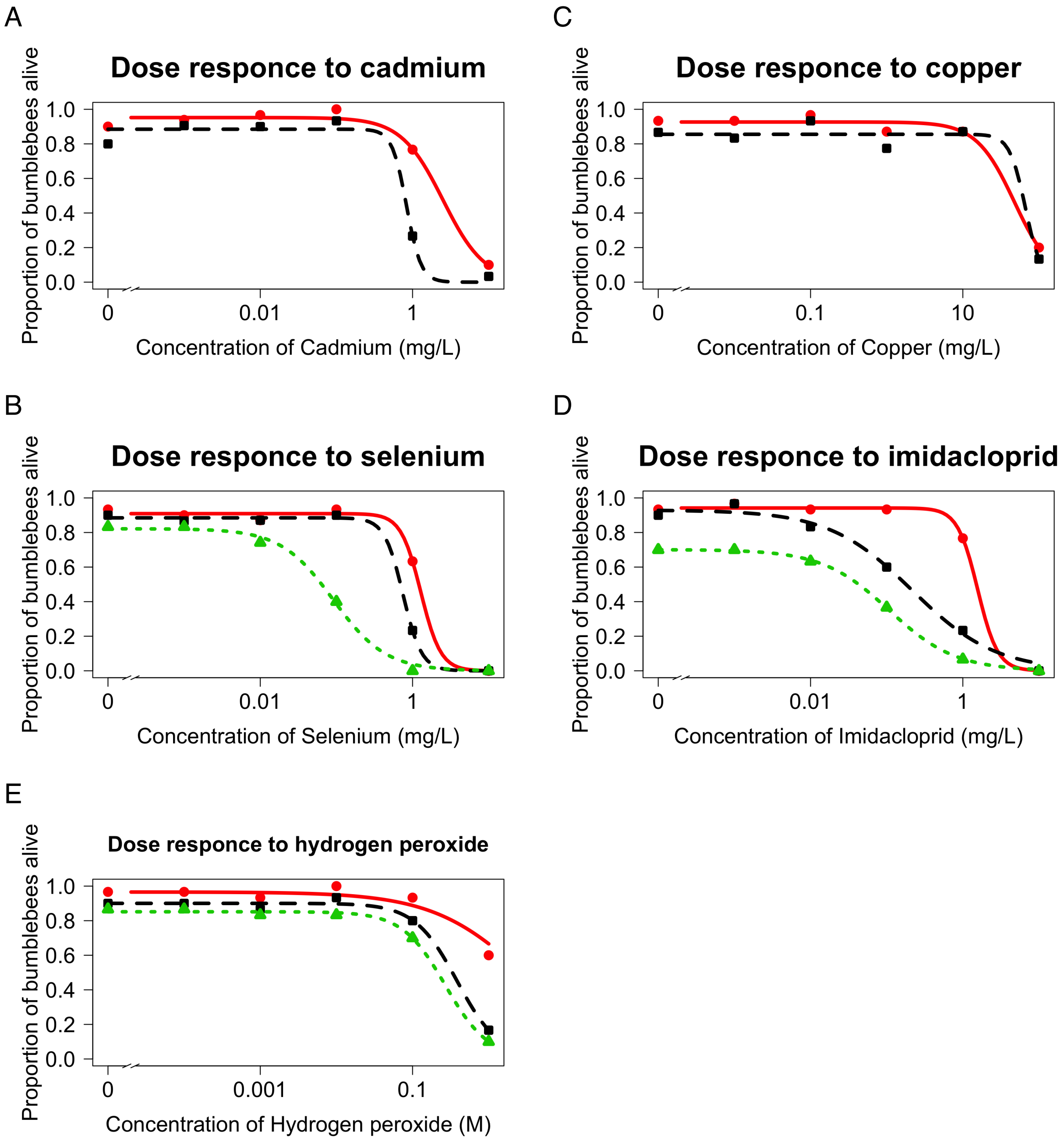


Supplemental Figure S3: Dose response curves for each toxicant treatment. Responses are shown based on 3 days (red solid line), 7 days (black dashed line), and 14 days (green dotted line).

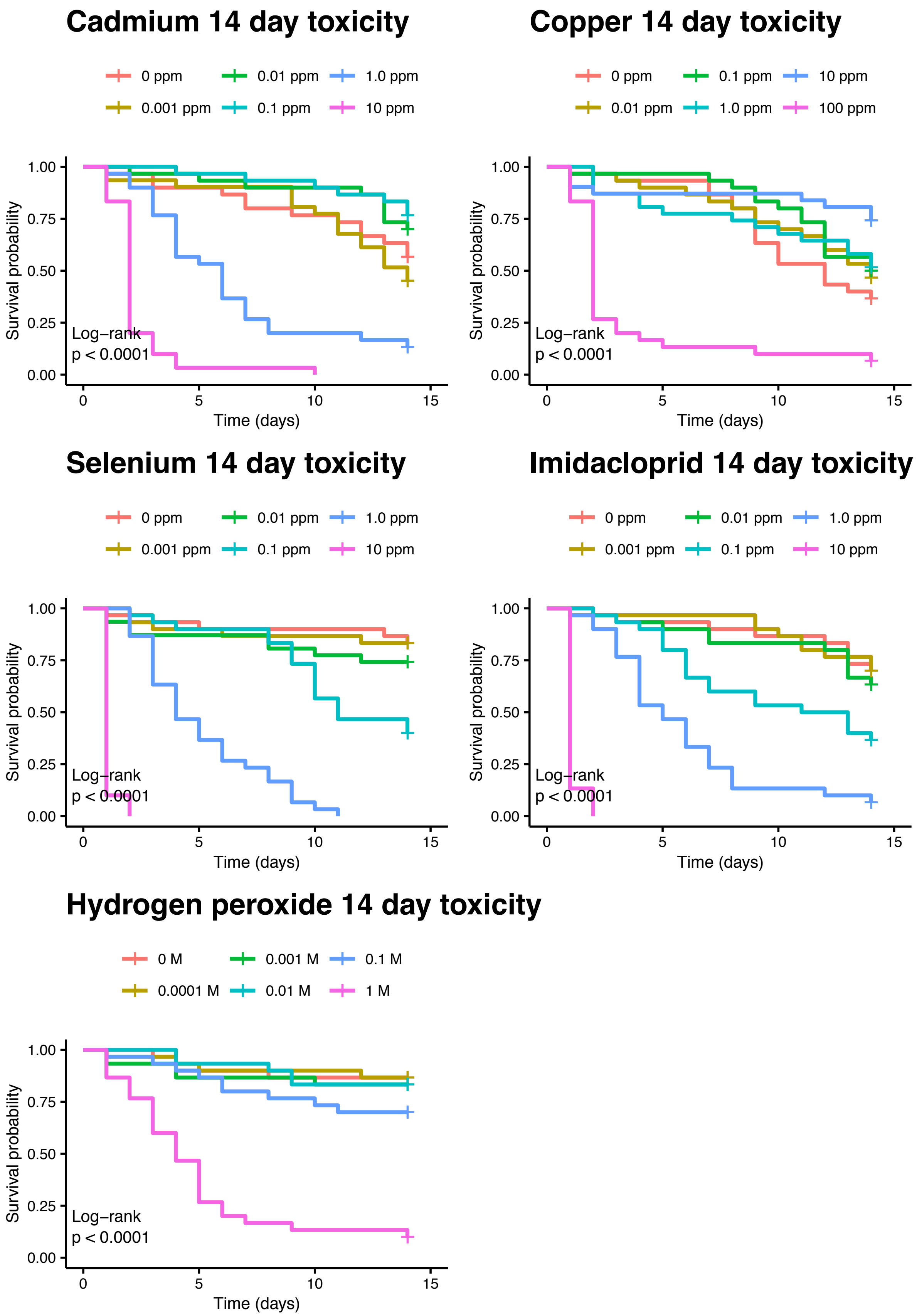
Cadmium 14 day toxicity
0 ppm
0.001 ppm
0.01 ppm
0.1 ppm
1.0 ppm
10 ppm
Survival probability
1.00
0.75
0.50
0.25
0.00
Log−rank
p < 0.0001
0
5
10
15
Time (days)
Copper 14 day toxicity
0 ppm
0.01 ppm
0.1 ppm
1.0 ppm
10 ppm
100 ppm
Survival probability
Log−rank
p < 0.0001
Time (days)
Selenium 14 day toxicity
0 ppm
0.001 ppm
0.01 ppm
0.1 ppm
1.0 ppm
10 ppm
Survival probability
Log−rank
p < 0.0001
Time (days)
Imidacloprid 14 day toxicity
0 ppm
0.001 ppm
0.01 ppm
0.1 ppm
1.0 ppm
10 ppm
Survival probability
Log−rank
p < 0.0001
Time (days)
Hydrogen peroxide 14 day toxicity
0 M
0.0001 M
0.001 M
0.01 M
0.1 M
1 M
Survival probability
Log−rank
p < 0.0001
Time (days)

Supplemental Figure S4: Survival probabilities of *Bombus impatiens* exposed to cadmium, copper, selenate, imidacloprid, and hydrogen peroxide for 14 days at each treatment dose. Toxicant exposure significantly reduced bee survivorship (cox proportional hazard test $P < 0.001$ for each compound).

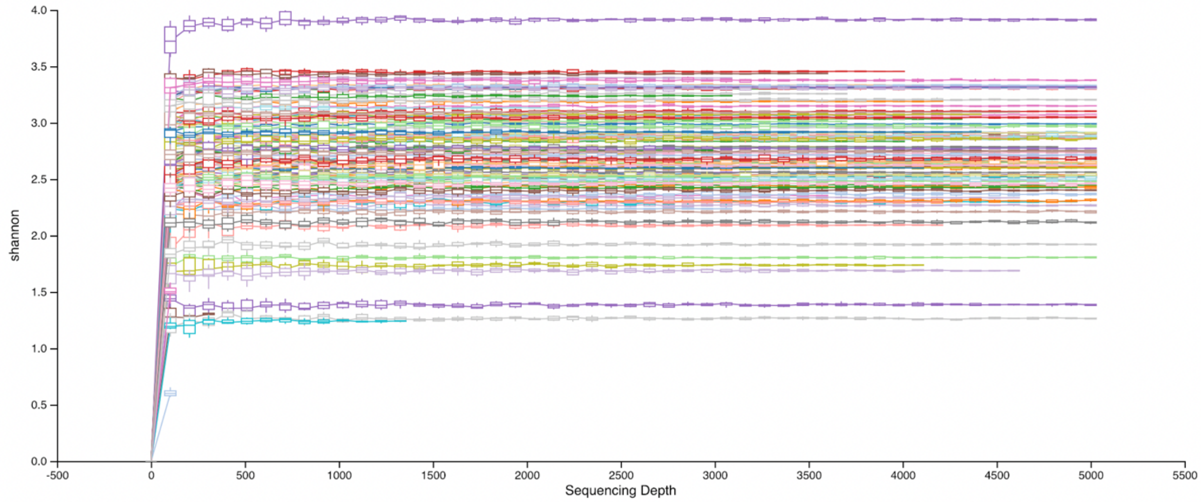


Supplemental Figure S5: Rarefaction analyses for each sample. Curves saturate at a sequencing depth under the read depth of 2,182 that we used for microbiome analyses.

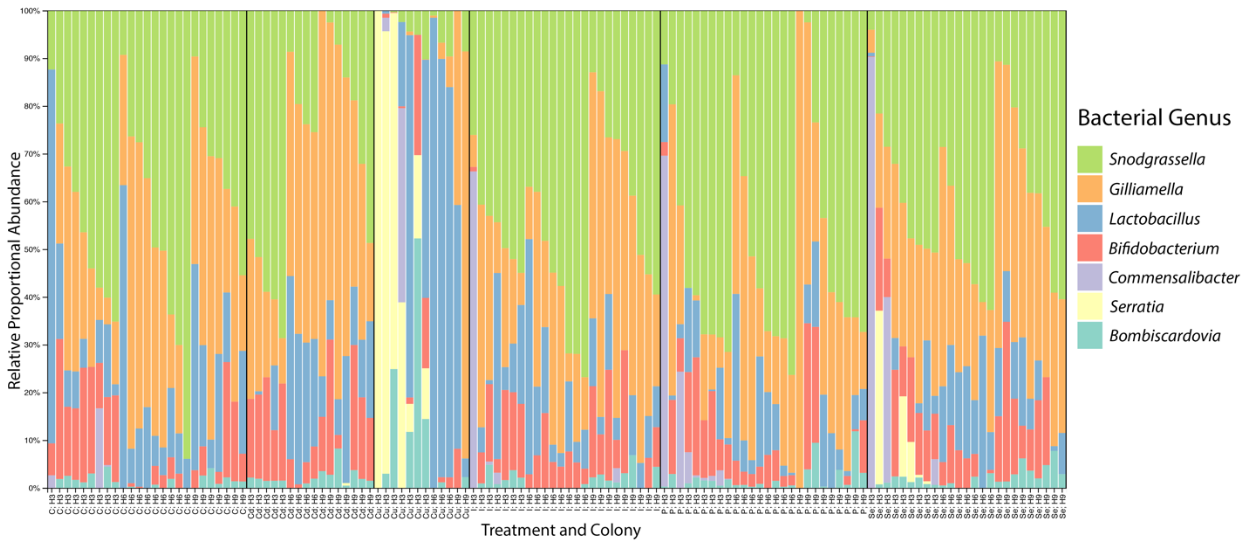


Supplemental Figure S6: Stacked bar plot showing bacterial genera that were present at greater than 1% abundance in each sample. Individual sample treatments are indicated by “C” for control, “Cd” for cadmium, “Cu” for copper, “I” for imidacloprid, “P” for hydrogen peroxide, and “Se” for selenate exposure. Colony of origin is denoted by H3, H6, or H9.

| Toxicant | Sample type | Mean Concentration (mg/kg) unless noted | Environment | Reference |
|---|---|---|---|---|
| Cadmium | *Helianthus annuus* leaves and flowers | 40 | Greenhouse | (1) |
| Cadmium | *Helianthus annuus* leaves and flowers | 161 | Greenhouse | (1) |
| Cadmium | *Solanum nigrum* leaves | 88 | Greenhouse | (2) |
| Cadmium | *Solanum melongena* leaves | 18 | Greenhouse | (2) |
| Cadmium | *Chamaecrista fasciculata* flowers | 1000 | Hydroponics | (3) |
| Cadmium | *Chamaecrista fasciculata* pollen | 4500 | Hydroponics | (3) |
| Cadmium | *Raphanus sativa* flowers | 13 | Greenhouse | (4) |
| Cadmium | Bee-collected pollen | 4 | Natural | (5) |
| Copper | *Helianthus annuus* leaves and flowers | 43 | Greenhouse | (1) |
| Copper | *Helianthus annuus* leaves and flowers | 75 | Greenhouse | (1) |
| Copper | *Raphanus sativa* flowers | 30 | Greenhouse | (4) |
| Hydrogen peroxide | *Cucurbita pepo* nectar | 0.096 mM | Garden | (6) |
| Hydrogen peroxide | *Nicotiana* sp. | 0.4 uMol/ul | Greenhouse | (7) |
| Hydrogen peroxide | *Nicotiana* sp. | 0.25 uMol/ul | Greenhouse | (7) |
| Hydrogen peroxide | *Nicotiana* sp. | 0.8 uMol/ul | Greenhouse | (7) |
| Hydrogen peroxide | *Nicotiana* sp. | 0.7 uMol/ul | Greenhouse | (7) |
| Hydrogen peroxide | *Nicotiana* sp. | 2.1 uMol/ul | Greenhouse | (7) |
| Hydrogen peroxide | *Nicotiana* sp. | 1.7 uMol/ul | Greenhouse | (7) |
| Hydrogen peroxide | *Nicotiana* sp. | 0.7 uMol/ul | Greenhouse | (7) |
| Hydrogen peroxide | *Nicotiana* sp. | 0.8 uMol/ul | Greenhouse | (7) |
| Hydrogen peroxide | *Nicotiana* sp. | 0.6 uMol/ul | Greenhouse | (7) |
| Imidacloprid | *Gossypium* sp. pollen | 9.03 | Agriculture | (8) |
| Imidacloprid | *Gossypium* sp. nectar | 0.38 | Agriculture | (8) |
| Imidacloprid | *Gossypium* sp. pollen | 33.1 | Agriculture | (8) |
| Imidacloprid | *Gossypium* sp. nectar | 0.87 | Agriculture | (8) |
| Imidacloprid | *Cucurbita pepo* pollen | 0.014 | Agriculture | (9) |
| Imidacloprid | *Cucurbita pepo* nectar | 0.01 | Agriculture | (9) |
| Imidacloprid | Bee-collected pollen | 0.012 | Agriculture | (10) |
| Imidacloprid | Bee-collected pollen | 39 | Agriculture | (11) |
| Selenium | *Stanleya pinnata* leaves | 1500 | Greenhouse | (12) |
| Selenium | *Stanleya pinnata* flowers | 3000 | Greenhouse | (12) |
| Selenium | *Brassica juncea* leaves | 375 | Greenhouse | (12) |
| Selenium | *Brassica juncea* flowers | 225 | Greenhouse | (12) |
| Selenium | *Astragalus bisulcatus* leaves | 6000 | Greenhouse | (13) |
| Selenium | *Astragalus bisulcatus* flowers | 8000 | Greenhouse | (13) |
| Selenium | *Stanleya pinnata* flowers | 1700 | Greenhouse | (13) |
| Selenium | *Stanleya pinnata* leaves | 1000 | Greenhouse | (13) |
| Selenium | *Astragalus bisulcatus* leaves | 3000 | Greenhouse | (14) |
| Selenium | *Astragalus bisulcatus* flowers | 4600 | Greenhouse | (14) |
| Selenium | *Astragalus bisulcatus* leaves | 2500 | Natural | (15) |
| Selenium | *Medicago sativa* | 20 | Natural | (15) |
| Selenium | *Stanleya pinnata* leaves | 1000 | Natural | (15) |
| Selenium | *Helianthus pumilus* leaves | 30 | Natural | (15) |
| Selenium | *Brassica juncea* pollen | 1700 | Greenhouse | (16) |
| Selenium | *Stanleya pinnata* pollen | 11000 | Greenhouse | (16) |
| Selenium | *Brassica juncea* nectar | 100 | Greenhouse | (16) |
| Selenium | *Stanleya pinnata* nectar | 150 | Greenhouse | (16) |

Table ST1: Concentrations of cadmium, copper, hydrogen peroxide, imidacloprid, and selenium found in various environments.

| Treatment | Dose | coef | exp(coef) | se(coef) | z | p |
|---|---|---|---|---|---|---|
| Cadmium | 0.001 mg/L | -0.7858 | 0.4558 | 0.7072 | -1.111 | 0.266522 |
| | 0.01 mg/L | -0.7494 | 0.4726 | 0.7071 | -1.06 | 0.289232 |
| | 0.1 mg/L | -1.199 | 0.3015 | 0.8165 | -1.468 | 0.142002 |
| | 1 mg/L | 1.7629 | 5.829 | 0.4659 | 3.784 | 0.000154 |
| | 10 mg/L | 3.5884 | 36.1748 | 0.4991 | 7.19 | 6.49E-13 |
| Copper | 0.01 mg/L | 0.26801 | 1.30736 | 0.67112 | 0.399 | 0.69 |
| | 0.1 mg/L | -0.70795 | 0.49265 | 0.86603 | -0.817 | 0.414 |
| | 1 mg/L | 0.63081 | 1.87913 | 0.6274 | 1.005 | 0.315 |
| | 10 mg/L | 0.03067 | 1.03114 | 0.70735 | 0.043 | 0.965 |
| | 100 mg/L | 2.7561 | 15.73828 | 0.55248 | 4.989 | 6.08E-07 |
| Selenate | 0.01 mg/L | 0.304063 | 1.355354 | 0.76378 | 0.398 | 0.6906 |
| | 0.1 mg/L | 0.320057 | 1.377207 | 0.763944 | 0.419 | 0.6752 |
| | 1 mg/L | -0.003469 | 0.996537 | 0.816549 | -0.004 | 0.9966 |
| | 10 mg/L | 2.46705 | 11.787624 | 0.618831 | 3.987 | 6.70E-05 |
| | 100 mg/L | 5.41444 | 224.626722 | 0.700112 | 7.734 | 1.04E-14 |
| Imidacloprid | 0.01 mg/L | -1.151 | 0.3163 | 1.1547 | -0.997 | 0.3189 |
| | 0.1 mg/L | 0.5323 | 1.7028 | 0.7304 | 0.729 | 0.4661 |
| | 1 mg/L | 1.5441 | 4.6836 | 0.646 | 2.39 | 0.0168 |
| | 10 mg/L | 2.6357 | 13.953 | 0.6179 | 4.265 | 2.00E-05 |
| | 100 mg/L | 6.0266 | 414.298 | 0.7428 | 8.113 | 4.94E-16 |
| Hydrogen Peroxide | 0.0001 M | -0.003538 | 0.996468 | 0.81666 | -0.004 | 0.997 |
| | 0.001 M | 0.276894 | 1.319027 | 0.775276 | 0.357 | 0.721 |
| | 0.01 M | -0.486411 | 0.614829 | 0.922243 | -0.527 | 0.598 |
| | 0.1 M | 0.712456 | 2.038993 | 0.707629 | 1.007 | 0.314 |
| | 1 M | 2.70872 | 15.010047 | 0.615737 | 4.399 | 1.09E-05 |

Supplemental Table ST2: Cox proportional hazard statistics for each treatment dose.

| | Shannon (α Diversity) | | Generalized Unifrac (β Diversity) | |
|---|---|---|---|---|
| Treatment Comparison | H | $P_{adj}$ | F | $P_{adj}$ |
| All Treatments | 24.21 | < 0.001 | 3.99 | < 0.001 |
| Cadmium vs Control | 0.04 | 0.851 | 2.37 | 0.017 |
| Copper vs Control | 2.85 | 0.137 | 7.86 | 0.002 |
| Imidacloprid vs Control | 3.61 | 0.115 | 0.48 | 0.960 |
| Peroxide vs Control | 4.61 | 0.079 | 3.54 | 0.002 |
| Selenate vs Control | 7.12 | 0.029 | 2.61 | 0.011 |

Supplemental Table ST3: Alpha and beta diversity statistics for the bacterial communities based on overall and pairwise treatments.

| **Treatment Comparison** | **ESV** | **$P_{adj}$** | **$Log_2$ Fold Change** | **ESV ID** |
|---|---|---|---|---|
| Cadmium vs Control | *Commensalibacter 1* | 0.026385 | -2.79877 | e7a9bec7a52e6d7a0b22163b009849b6 |
| Copper vs Control | *Serratia 1* | 1.86E-56 | 11.46145 | 5e665e4087d910f92acdda1a4af8ffd6 |
| | *Serratia 2* | 1.16E-55 | 11.59668 | 9fc456429ad9bbf498f75dec8619dcd9 |
| | *Gilliamella 1* | 7.32E-16 | -4.79262 | 3723f20958224625ccef8bce3a3d827a |
| | *Gilliamella 2* | 5.46E-12 | -4.39918 | 8e59691bc792d8c8d11b406924320a3c |
| | *Bombiscardovia 1* | 4.02E-11 | 5.23348 | 996f16a5ff6b3e6ee56c13c544f3f2e3 |
| | *Snodgrassella 1* | 7.29E-10 | -2.91227 | b776638f1aa719468743535ae2ea66c2 |
| | *Snodgrassella 2* | 7.29E-10 | -3.03113 | f031d808057dac4dce64a9055f3cec01 |
| | *Bombiscardovia 2* | 1.46E-08 | 4.602601 | aa33497d23e80f30542f4cf63a67d936 |
| | *Commensalibacter 1* | 3.72E-06 | 4.090161 | 07254eace7b0e961855777ccc842471b |
| | *Gilliamella 3* | 0.002848 | 2.969599 | 445175fd127131bf3aca48c508278bc0 |
| | *Gilliamella 4* | 0.013032 | 2.4478 | 8f0ceaec8c70d52867b56a2437817325 |
| | *Lactobacillus 1* | 0.016473 | 1.925286 | 1b942649ef3eab598cd77cd0d2f7bfb4 |
| | *Lactobacillus 2* | 0.025571 | 1.769691 | fe3188b7068f70f5f395682b98ab9b7c |
| Imidacloprid vs Control | No significant differences found. | N/A | N/A | N/A |
| Peroxide vs Control | *Commensalibacter 1* | 0.009931 | 2.556807 | 07254eace7b0e961855777ccc842471b |
| Selenate vs Control | *Commensalibacter 1* | 1.27E-09 | 4.711861 | 07254eace7b0e961855777ccc842471b |
| | *Commensalibacter 1* | 0.002314 | 2.530897 | e7a9bec7a52e6d7a0b22163b009849b6 |
| | *Lactobacillus 2* | 0.006638 | -2.03093 | fe3188b7068f70f5f395682b98ab9b7c |
| | *Lactobacillus 1* | 0.01297 | -1.86744 | 1b942649ef3eab598cd77cd0d2f7bfb4 |
| | *Snodgrassella 1* | 0.014975 | -1.04744 | b776638f1aa719468743535ae2ea66c2 |
| | *Gilliamella 1* | 0.014975 | -1.30594 | 3723f20958224625ccef8bce3a3d827a |
| | *Gilliamella 2* | 0.016117 | -1.36575 | 8e59691bc792d8c8d11b406924320a3c |
| | *Snodgrassella 2* | 0.028255 | -0.96929 | f031d808057dac4dce64a9055f3cec01 |

Supplemental Table ST4: Statistics and ESV IDs associated with significantly differentially abundant bacterial ESVs between treatments.

Supplemental File SF1: Bacterial strain ID and NCBI accession numbers for genomes used in this study.

Supplemental File SF2: Exact sequence variant (ESV) table with ESV counts per sample, SILVA taxonomy, and the top BLAST hit for each ESV.

Supplemental References:


1. Dhiman SS, Zhao X, Li J, Kim D, Kalia VC, Kim I-W, et al. Metal accumulation by sunflower (*Helianthus annuus* L.) and the efficacy of its biomass in enzymatic saccharification. Kim KH, editor. PLoS One. 2017 Apr 24;12(4):e0175845. Available from: 10.1371/journal.pone.0175845
2. Sun RL, Zhou QX, Jin CX. Cadmium accumulation in relation to organic acids in leaves of *Solanum nigrum* L. as a newly found cadmium hyperaccumulator. Plant Soil. 2006 Jul;285(1–2):125–34. Available from: 10.1007/s11104-006-0064-6
3. Henson TM, Cory W, Rutter MT. Extensive variation in cadmium tolerance and accumulation among populations of *Chamaecrista fasciculata*. PLoS One. 2013;8(5):e63200. Available from: 10.1371/journal.pone.0063200
4. Hladun KR, Parker DR, Trumble JT. Cadmium, copper, and lead accumulation and bioconcentration in the vegetative and reproductive organs of *Raphanus sativus*: implications for plant performance and pollination. J Chem Ecol. 2015 Apr 7;41(4):386–95. Available from: 10.1007/s10886-015-0569-7
5. Conti ME, Botrè F. Honeybees and their products as potential bioindicators of heavy metals contamination. Environ Monit Assess. 2001; Available from: 10.1023/A:1010719107006
6. Nocentini D, Guarnieri M, Soligo C. Nectar defense and hydrogen peroxide in floral nectar of *Cucurbita pep*o. Acta Agrobot. 2015;68(2):187–93. Available from: 10.5586/aa.2015.009
7. Silva FA, Guirgis A, Thornburg R. nectar analysis throughout the genus *Nicotiana* suggests conserved mechanisms of nectar production and biochemical action. Front Plant Sci. 2018 Jul 30;9. Available from: 10.3389/fpls.2018.01100
8. Jiang J, Ma D, Zou N, Yu X, Zhang Z, Liu F, et al. Concentrations of imidacloprid and thiamethoxam in pollen, nectar and leaves from seed-dressed cotton crops and their potential risk to honeybees (*Apis mellifera* L.). Chemosphere. 2018 Jun 1;201:159–67. Available from: 10.1016/j.chemosphere.2018.02.168
9. Stoner KA, Eitzer BD. Movement of soil-applied imidacloprid and thiamethoxam into nectar and pollen of squash (*Cucurbita pepo*). PLoS One. 2012 Jun 27;7(6):e39114. Available from: 10.1371/journal.pone.0039114
10. Chauzat MP, Faucon JP, Martel AC, Lachaize J, Cougoule N, Aubert M. A survey of pesticide residues in pollen loads collected by honey bees in France. Vol. 99, Journal of Economic Entomology. Entomological Society of America; 2006. p. 253–62. Available from: 10.1093/jee/99.2.253
11. Mullin CA, Frazier M, Frazier JL, Ashcraft S, Simonds R, vanEngelsdorp D, et al. High Levels of miticides and agrochemicals in North American apiaries: implications for honey bee health. PLoS One. 2010 Mar 19;5(3):e9754. Available from: 10.1371/journal.pone.0009754
12. Quinn CF, Prins CN, Freeman JL, Gross AM, Hantzis LJ, Reynolds RJB, et al. Selenium accumulation in flowers and its effects on pollination. New Phytol. 2011 Nov 1;192(3):727–37. Available from: 10.1111/j.1469-8137.2011.03832.x
13. Freeman JL, Zhang LH, Marcus MA, Fakra S, McGrath SP, Pilon-Smits EAH. Spatial imaging, speciation, and quantification of selenium in the hyperaccumulator plants *Astragalus bisulcatus* and *Stanleya pinnat*a. Plant Physiol. 2006 Sep;142(1):124–34.

Available from: 10.1104/pp.106.081158
14. Valdez Barillas JR, Quinn CF, Freeman JL, Lindblom SD, Fakra SC, Marcus MA, et al. Selenium distribution and speciation in the hyperaccumulator *Astragalus bisulcatus* and associated ecological partners. Plant Physiol. 2012 Aug;159(4):1834–44. Available from: 10.1104/pp.112.199307
15. Galeas ML, Klamper EM, Bennett LE, Freeman JL, Kondratieff BC, Quinn CF, et al. Selenium hyperaccumulation reduces plant arthropod loads in the field. New Phytol. 2008 Feb;177(3):715–24. Available from: 10.1111/j.1469-8137.2007.02285.x
16. Hladun KR, Parker DR, Trumble JT. Selenium accumulation in the floral tissues of two Brassicaceae species and its impact on floral traits and plant performance. Environ Exp Bot. 2011;74(0):90–7. Available from: 10.1016/j.envexpbot.2011.05.003
